# Orally surface engineered probiotic system for ulcerative colitis therapy *via* modulating gut microbiota and immune homeostasis

**DOI:** 10.1101/2024.08.22.609086

**Authors:** Junyu Liu, Zhihao Fang, Xiaopeng Zhang, Yun Chen, Fujia Kou, Xiaobin Li, Long Huang, Huanxian Yang, Yilin Zheng, Yuqing Huang, Yi Wang, Xueqing Qiu, Jun Ge, Yong Qian, Xin-Hui Xing, Can Yang Zhang

## Abstract

Oral probiotics have emerged as a promising therapeutical strategy for effectively managing ulcerative colitis (UC) in the world. The existing researches successfully preserve probiotic viability in the upper gastrointestinal tract, but they fall short in achieving precise release profile and effective colonization of probiotics at the site of colitis in colon as well as the understanding of therapeutic mechanism with suboptimal efficacy. This constraint poses a significant barrier to rational design and effective development of oral probiotic system. Here, we fabricate an orally layered-by-layered probiotic (*Escherichia coli* Nissle 1917, *EcN*) system (so-called CML@*EcN*) based on carboxymethyl modified lignin (CML) which can significantly protect *EcN* in gastric and small intestinal microenvironment and efficiently control release *EcN* in colon. Interestingly, the release and proliferation process of *EcN* in the colon are detected and can be modeled as a plug-flow mode. We derive a mathematical expression that highly matches the experimental results, providing a theoretical basis for quantitatively calculation of the release process. Furthermore, CML@*EcN* significantly alleviates symptoms in dextran sulfate sodium (DSS)-induced UC mice by modulating the multiple organ immune disorder and gut microbiota (GM)/metabolites profile as well as reconstructing the colon barrier. Importantly, we explore the mechanism of immune homeostasis regulation *via* metabolism from GM. Moreover, the CML armour is excreted and would not influence the process. This work not only reports a novel oral therapeutic for UC treatment with high efficacy as well as the exploration of mechanism between GM and immune homeostasis, but also provides a general approach to engineer probiotics for oral administration.

## Main

Gut microbiota (GM) dysbiosis and immune dyshomeostasis are pivotal in the pathogenesis of ulcerative colitis (UC). This manifests predominantly through the infiltration of immune cells into the gut lumen and the aberrant translocation of GM into body tissues^1^. Such disruption leads to a cascade of immune dysregulation affecting multiple organ systems, notably the intestines and spleen. The ensuing immunological perturbation not only amplifies the impairment of the intestinal barrier but also escalates the pathological progression of UC^2, 3^. Existing anti-inflammatory or immunosuppressive agents have limited efficacy due to the singular mechanism of action along with obvious side effects^4, 5^. Specific probiotics, by enhancing GM composition and repairing the gut barrier with minimal side effects, have emerged as a promising therapeutic approach for UC^6, 7^. By far, *Escherichia coli* Nissle 1917 (*EcN*) still being the only registered probiotic drug for UC therapy. However, current formulations of *EcN* require a daily oral dose of nearly 10^10^ colony forming unit (CFU), yet the efficacy remains unsatisfactory. This is primarily due to the inability to achieve effective colonization of *EcN* at site of UC^8^.

Oral delivery systems with enhanced colonization of probiotics at focal site and reduced side effect have been recognized as a promising strategy for UC treatment^9, 10^. For instance, encapsulation in various membranes or liposomes can maintain high probiotic activity within the digestive tract, and surface modification of probiotics can improve the permeability of the delivery system through the intestinal mucus layer, thereby enhancing intestinal colonization rates^11–13^. Nevertheless, the lesions of UC are primarily located in colon. Existing oral delivery systems commonly result in the premature release of probiotics in the small intestine, which limits their colonization efficacy in colon and impedes research into the therapeutic mechanisms of probiotics for UC. Therefore, there is an urgent need to develop innovative oral delivery systems that can achieve efficient *in situ* colonization of probiotics in the colon, enabling effective treatment of UC and elucidation of the underlying mechanisms.

Lignin widespread in plants is resistant to digestion and absorption by mammals with high biosafety^14^. To improve the solubility and stimuli-responsibility of lignin, the hydroxyl residues were transformed as sodium carboxymethyl (Supplementary Fig. 1), resulting in carboxymethyl-modified lignin (CML) in this work. The CML was used as a suit of armour for *EcN* to prepare oral probiotic system (CML@*EcN*) *via* layer-by-layer surface engineered process, which can precisely and effectively deliver *EcN* to the colitis site, followed by modulating the GM and regulating the immune homeostasis. Furthermore, we carefully explore the delivery mechanism *in vivo* and *in vitro* basing on the biological experiments and theoretical analysis using a plug-flow model. We next carefully reveal the therapeutic mechanism based on synergistic effect of modulation of GM and immune homeostasis, meanwhile, evaluate the therapeutic efficacy (Fig. 1). Given the universality of the delivery system and kinetic model constructed in this work, we anticipate that it can be used to the treatment of colon diseases by orally administered probiotics *via* modulating GM and immune homeostasis.

**Fig. 1.**
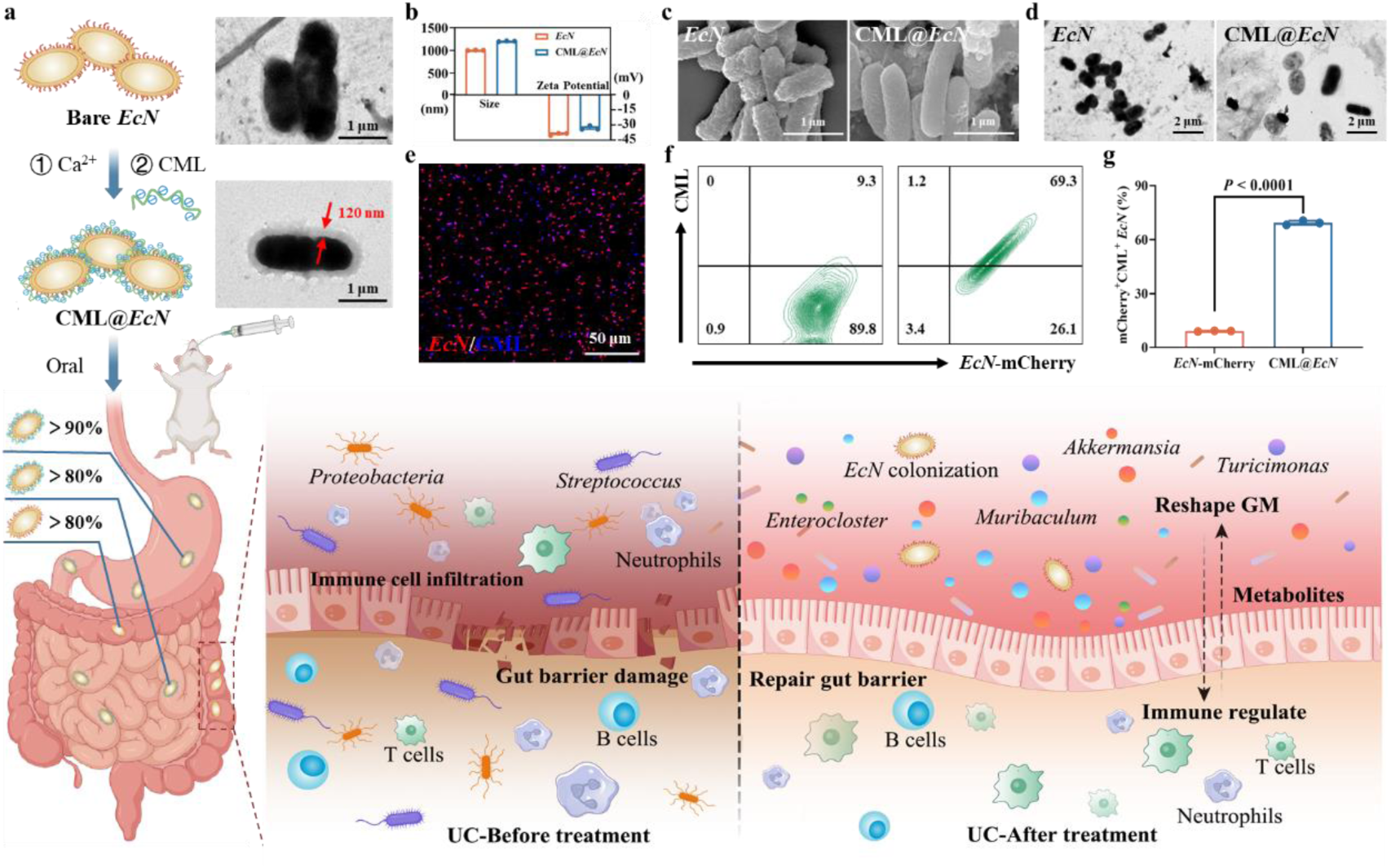
Development of oral probiotic delivery system for UC management. **a,** Schematic of the oral administration of CML@*EcN* for UC treatment via the colon barrier repair, GM reconstitution and immune homeostasis. CML@*EcN* was prepared by the method of electrostatic adsorption, and achieves the stabilization of *EcN* in the stomach and small intestine, as well as controlled release and colonization in the colon. Furthermore, CML@*EcN* co-regulated the immune system function and gut environment of UC mice, thereby restoring gut barrier function and achieving therapeutic effects for UC. **b,** DLS is used to monitor changes in particle size and Zeta-potential before and after encapsulate. **c,** SEM (scale bar, 1 μm) & **d,** TEM (scale bar, 2 μm) were used to observe the morphology of the system. **e,** CLSM were used to visual characterization of the mCherry labeled EcN system, scale bar, 50 μm. **f,g,** flow cytometry results were utilized to confirmed the successful construction of the encapsulation system.

### Preparation of CML@*EcN*

The hydroxyl residues in lignin were transformed as carboxymethyl ones to enhance the solubility and introduce the pH-sensitivity (Supplementary Fig. 1a), resulting in armour in this work. Fourier Transform Infrared Spectroscopy (FTIR) was utilized to confirm the successful modification of lignin (Supplementary Fig. 1b). The disappearance of the C-Cl characteristic peak at 776 cm⁻¹and the emergence of a new benzene ring para-substitution peak at 835 cm⁻¹ in spectrum of CML proved the successful carboxymethylation of lignin (named CML). Additionally, the proccess conditions like reaction temperature and feed ratio were optimized (Supplementary Table 1). Aqueous potentiometric titration results demonstrated that the carboxyl content of the optimized CML could reach 3.2439 mmol/g, and the zeta-potential was measured as −31.83 mV (Supplementary Table 2).

*In vivo* and *in vitro* biosafety of CML was validated immediately. NCM-460 cells were used to evaluate the biosafety *in vitro*, showing that the cell viability could maintain over 90% after 48 h of incubation even at the highest concentration of 300 μg/mL of CML (Supplementary Fig. 2), which indicated the negligible cytotoxicity. Importantly, C57 BL/6N mice were used to evaluate the biosafety of CML *in vivo*. After 15 days of continuous gavage, CML (both low does and high dose) did not cause significant weight changes in mice (Supplementary Fig. 3a). No abnormalities in blood biochemical indicators related to mouse liver function, kidney function, and inorganic ions were observed (Supplementary Fig. 3b-d). Additionally, the major organs (intestine, heart, liver, spleen, lung and kidney) showed no abnormal histological characteristics (Supplementary Fig. 4). Taken together, all these findings indicated that the CML had high biosafety, showing the huge potential in biomedical application.

Then, CML@*EcN* was fabricated by layer-by-layer technique *via* electrostatic interaction (Supplementary Fig. 5). The fabrication process, including the feed ratio, incubation time of Ca^2+^ and CML, was successfully optimized (Supplementary Fig. S6). The CML@*EcN* system prepared at concentration of Ca^2+^ and CML at 1.0 M and 0.8 g/L with incubation time of 30 min and 60 min, respectively, was selected as sample for the following studies. After that, the physicochemical properties were carefully characterized. The particle size of the CML@*EcN* slightly increased from approximately 1000 nm to 1200 nm (Fig. 1a,b) and the zeta-potential also exhibited surface charge reversal (Fig. 1b) measured by dynamic light scattering (DLS), confirming the successful fabrication of CML@*EcN* system. Moreover, scanning electron microscopy (SEM) and transmission electron microscopy (TEM) were utilized to observe the morphology of *EcN* before and after encapsulation. The results revealed that the surface of the CML@*EcN* appeared noticeably smoother than that of the unwrapped *EcN* (Fig. 1c). Additionally, a film with a thickness of approximately 120 nm was prominently observed on the surface of the encapsulated *EcN* (Fig. 1d). Subsequently, mCherry-labeled *EcN* was used for visual characterization of the delivery system. Confocal laser scanning microscopy (CLSM) images demonstrated significant overlap between *EcN* and CML (inherent blue fluorescence) signals, indicating successful encapsulation of the majority of *EcN* by CML (Fig. 1e, Supplementary Fig. 7). Flow cytometry quantification results further confirmed a fabrication success rate of 69.3% for CML@*EcN* (Fig. 1f,g). These characterizations collectively proved the successful fabrication and encapsulation of *EcN*.

### *In vitro* simulation of CML@*EcN* existence in the digestive tract

The stability of CML@*EcN* system in gastric acid was evaluated through an *in vitro* simulated gastric environment experiment. The results demonstrated that unwrapped *EcN* was severely inactivated after 30 min of exposure to the simulated gastric environment. In contrast, CML@*EcN* was able to maintain *EcN* activity at levels exceeding 90% for at least 2 h under the same conditions (Fig. 2a,b). Additionally, their morphologies were confirmed using TEM, showing that *EcN* exposed to the simulated gastric environment was largely degraded, whereas CML@*EcN* maintained the normal morphology of *EcN* (Fig. 2c). Furthermore, live/dead bacterial staining was employed for qualitative and quantitative analysis of *EcN* survival in the simulated gastric environment using CLSM and flow cytomentry. The results indicated that CML@*EcN* effectively protected *EcN* with higher than 90% survival in this environment (Fig. 2d,f, Supplementary Fig. 8). At pH of 1.2 (simulating the gastric environment) and pH 6.8 (simulating the intestinal environment), *EcN* exhibited minimal release with less than 30% after incubation of 8 h. However, at pH of 7.4 (simulating the colonic environment), over 85% of *EcN* was released (Supplementary Fig. 9). The different release profiles of *EcN* could be induced by the carboxyl residues in CML, which could be ionization/deionization at different pH conditions. These findings suggested that CML@*EcN* can remain encapsulated while passing through the stomach and small intestine, thereby ensuring the release of highly active *EcN* in the colon.

**Fig. 2.**
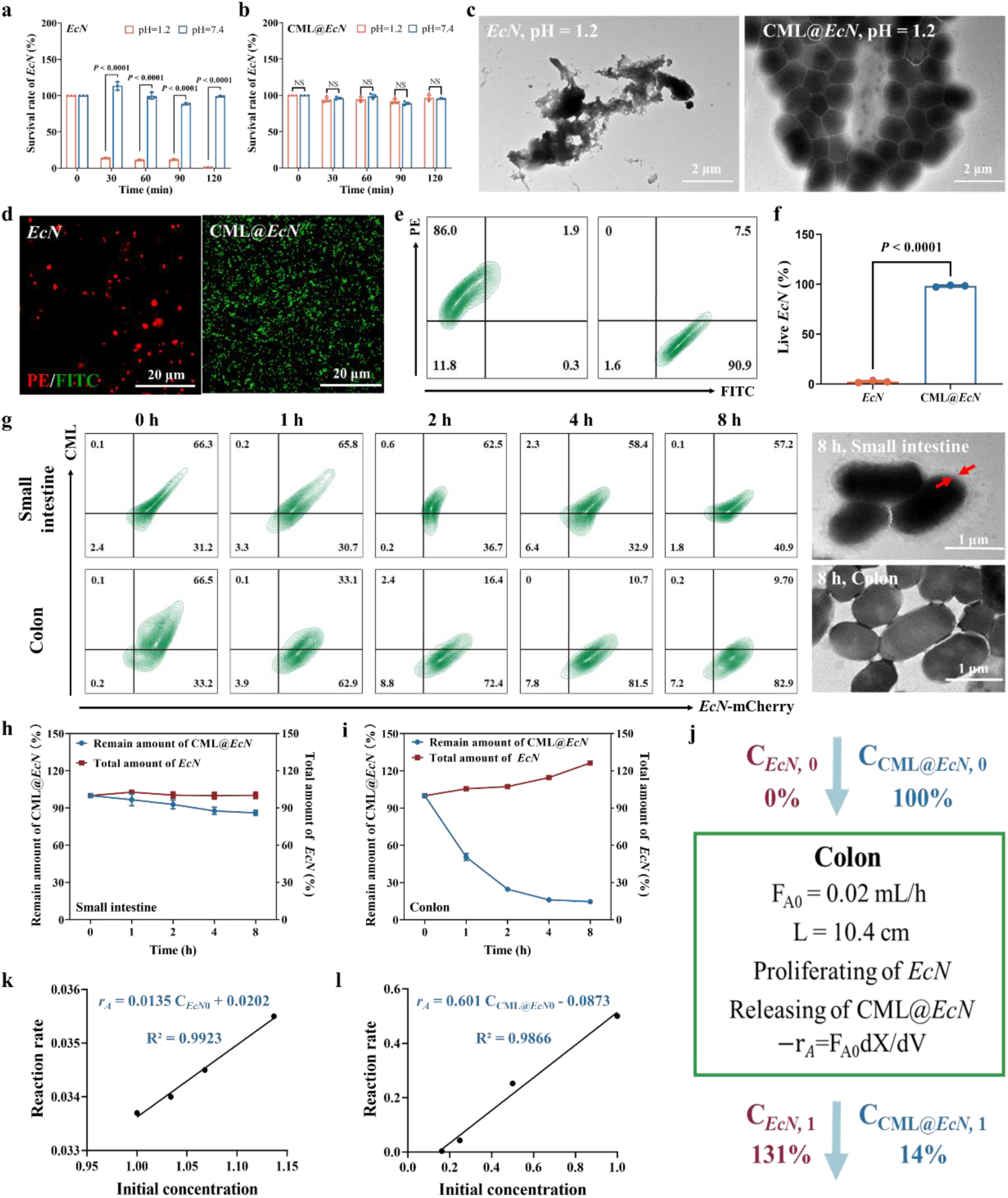
*In vitro* study on the existence form of CML@*EcN* in various segments of the digestive tract. **a,b,** *EcN* and CML@*EcN* survival rate within 2 h under simulated stomach environment and a pH neutral environment, respectively. **c,** TEM was employed to observe the morphology of *EcN* and CML@*EcN* in simulated gastric fluid after a 2 h incubation. **d,** CLSM was used to observe *EcN* and CML@*EcN* after viability staining. Scale bar, 20 μm. **e,f,** flow cytometry were used to quantitatively evaluated the activity of *EcN* under simulated gastric environment *in vitro*. Furthermore, mCherry labeled *EcN* were used to fabricate CML@*EcN*, and the systems were placed in simulated small intestine and colon environment, respectively. **g,** Flow cytometry was used to detect with samples taken at different time points. **g,** TEM was utilized to observe the morphological characteristics of CML@*EcN* after 8 h incubation in simulated intestinal and colonic fluids. Scale bar, 1 μm. **h,i,** The total amount of EcN and the degradation ratio of CML@*EcN* were presented in line graphs. **j,** The plug-flow model is used to simulate the release process of CML@*EcN* in the colon. **k,l,** The fitted release curve and proliferation curve of CML@*EcN* in the colon.

To further investigate the stability and release behavior of CML@*EcN*, mCherry-labeled *EcN* was utilized to fabricate CML@*EcN*-mCherry, which was incubated in simulated intestinal fluid and colonic fluid. At various incubation time points, samples were extracted and quantitatively analyzed using flow cytometry. The results showed that over 80% of CML@*EcN* remained stable in the simulated intestinal fluid. Conversely, in the simulated colonic fluid, over 80% of CML@*EcN* degraded within 2 h, resulting in the release of *EcN* (Fig. 2g). The morphology of samples was visually confirmed by TEM. After 8 h of incubation, most *EcN* in the simulated small intestine fluid still had a clear membrane visible on the surface (armor belt), whereas the membrane on the surface of *EcN* in the simulated colon fluid has disappeared (armor remover). Additionally, release curves for CML@*EcN* and proliferation curves for *EcN* in the small intestine and colon environments were plotted, respectively (Fig. 2h,i). These curves further illustrated the differential stability and release kinetics of CML@*EcN* in response to the distinct pH environments, supporting the targeted release mechanism of the delivery system.

To explore the release and proliferation process of CML@*EcN* in the gastrointestinal tract universally, we aimed to establish a dynamic simulation model. The pH and enzyme composition of each segment of the digestive tract are relatively stable, allowing us to approximate the intestine as a tubular reactor. Given that most *EcN* is released in the colon, a plug-flow model is suitable for simulating the release and proliferation process of CML@*EcN* in this section (Fig. 2j). Based on the average length, cross-sectional area, and residence time of food in the colon of mice^15^, the concentration-reaction rate curve for CML@*EcN* was established. The fitting results indicated that the release and proliferation processes of *EcN* in the colon can be approximated as first-order reactions (Fig. 2k,l). The results for the system in small intestine were showed in Supplementary Fig. 10. According to the derived formula (Showed in supplementary information), the half-release time of *EcN* was approximately 1.46 hours, with around 86% of *EcN* released in the colon, aligning closely with experimental results (Fig. 2g-i). We further extended our predictions to the colonization behavior of *EcN* in the colon. Assuming that 100% of CML@*EcN* enters the colon, after 20 hours (the typical residence time of substances in the colon), the total abundance of *EcN* was projected to reach 145%. This comprises 14% as unreleased CML@*EcN*, 86% as released *EcN*, and 45% as newly proliferated *EcN*, resulting in free *EcN* (131%) colonizing the colon, which was also confirmed by experiments (Fig. 2i). The predicted final colonization of *EcN* was estimated to represent approximately 0.1% of the total gut microbiota (GM) abundance. This prediction was validated by subsequent *in vivo* distribution experiments (Fig. 3), demonstrating that this model is suitable to simulate the release and proliferation processes of similar colon-targeted oral probiotic preparations. This validation underscores the robustness of the plug-flow model and its predictive power, providing a reliable tool for future research and development of targeted probiotic delivery systems (supporting information for the derivation process).

**Fig. 3.**
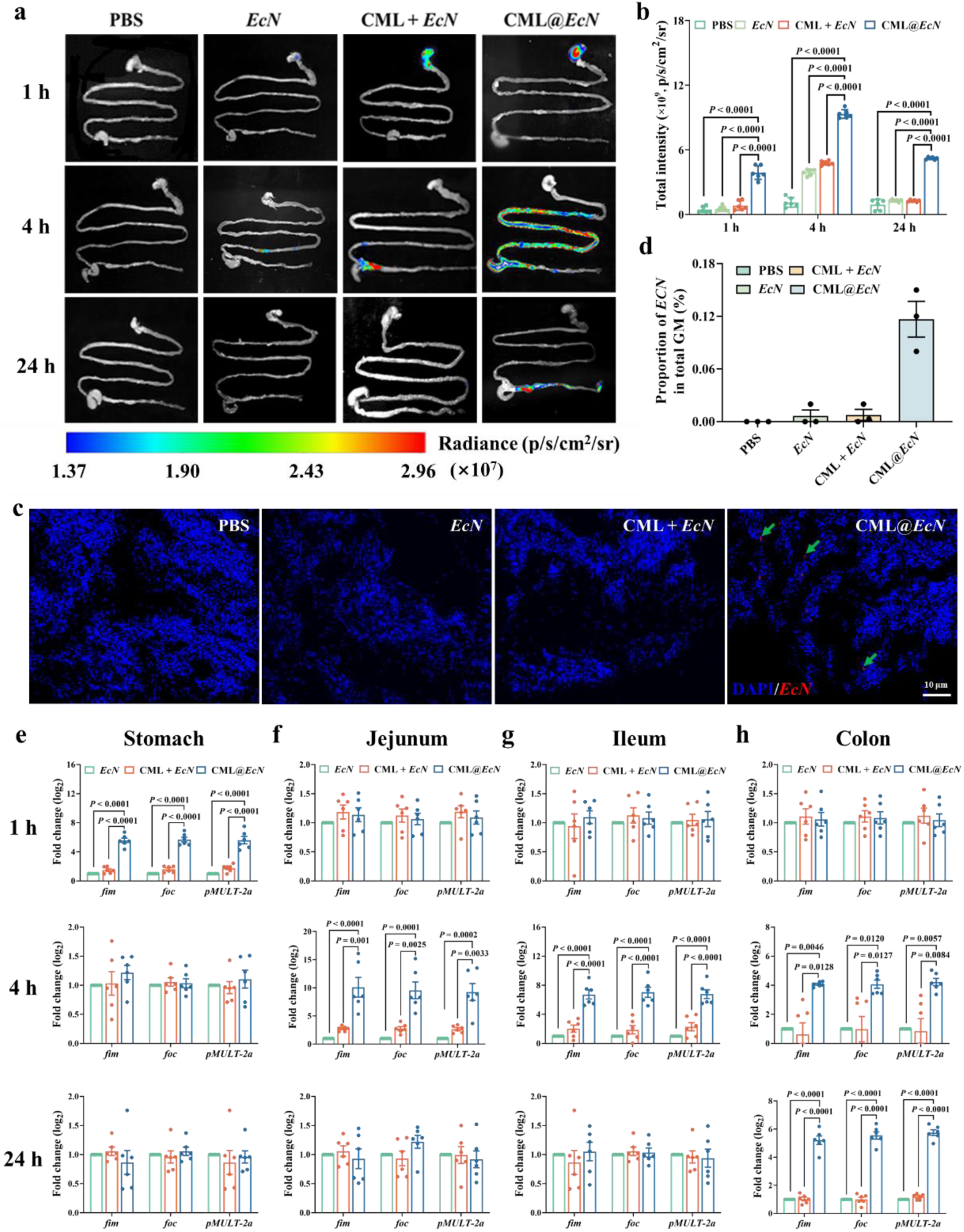
Qualitative and quantitative analysis of the *in vivo* biodistribution of different oral *EcN* formulations. **a,b,** IVIS was used to evaluate the biodistribution of oral *EcN* at 1 h, 4 h and 24 h, respectively. Indicating CML@*EcN* significantly improved the survival and colonization ability of *EcN* in the entire intestine of mice within 24 hours. **c,d,** 24h colon contents of each group were collected for FISH detection (arrows point to the successfully colonized single *EcN*). Scale bar, 10 μm. **e-h,** RT-PCR was used to quantitatively analyze the content of *EcN* at 1 h, 4 h and 24 h of different segments of the digestive tract, which quantitatively demonstrated the results of IVIS.

Taken together, it can be inferred that CML@*EcN* can remain stable in the stomach and small intestine, while achieving controlled release on-demand and proliferation in the colon region.

### *In vivo* biodistribution of CML@*EcN*

To further investigate the distribution of *EcN in vivo* after oral administration, *In vivo* imaging system (IVIS) and RT-PCR were used to qualitatively and quantitatively analyze *EcN* in various segments of the digestive tract, respectively. The IVIS results indicated that mice orally administered with CML@*EcN* showed specific signal enrichment in the stomach at 1 h, significant signal enhancement in the intestine at 4 h, and specific enrichment in the colon at 24 h compared to free *EcN* and the physical mixture of CML and *EcN* (CMI+*EcN*) group (Fig. 3a,b, Supplementary Fig.11a). No signal distribution was observed in other important organs of each group in mice (Supplementary Fig. 11b,c). These results indicated that CML@*EcN* effectively protected the activity of *EcN* in the stomach and intestines, followed by achieving colonization of *EcN* in the colon within 24 h. Furthermore, fluorescence *in situ* hybridization (FISH) of colonic contents samples at 24 h visually confirmed the successful colonization of *EcN* in the colon region (Fig. 3c,d, Supplementary Fig. 12). The quantitative analysis of *EcN* content in different segments of the gastrointestinal tract corroborated this phenomenon. Importantly, quantitative results of FISH showed that *EcN* accounted for 0.12% of the GM, which is highly consistent with the previous results calculated based on the plug-flow model (0.1%). Furthermore, RT-PCR results showed that after 24 h of gavage, CML@*EcN* significantly increased the *EcN* content in the colon of UC mice (approximately 2^6^ times higher compared to the other groups) (Fig. 3e-g), and *EcN* could still be detected in the colonic contents after 72 hours of gavage (Supplementary Fig. 13). Additionally, following the release of *EcN*, almost 70% of CML was excreted with the faeces (Supplementary Fig. 14), indicating that CML does not accumulate in the body or participate in organism metabolism, posing no biosafety hazards as confirmed as forementioned.

Indeed, the *in vivo* biodistribution results further validate the applicability of the kinetic model and formulas used in the study. These findings confirm that CML@*EcN* successfully protects the activity of *EcN* in the stomach and small intestine, allowing for release on-demand and colonization specifically in the colon region. The effective colonization of *EcN* in the colon is of paramount importance for enhancing the efficacy of *EcN*-based therapies and facilitating subsequent mechanism research.

### Remission of UC *in vivo* and gut barrier repair effect of CML@*EcN*

DSS-induced UC-bearing mouse model was used to evaluate the therapeutic efficacy of CML@*EcN* (Fig. 4a). After 7 days of treatment, UC-bearing mice without treatment showed significant weight loss (Fig. 4b), increased spleen weight index (Fig. 4c) and shortened colon length (Fig. 4d,e), which are typical UC symptoms, while the intervention of CML@*EcN* obviously alleviated the above UC phenotypes, proving the high therapeutic efficacy. Moreover, mice treated with CML@*EcN* showed significantly better effect in improvement UC phenotype than those in the delivery material mixed directly and intervened free *EcN* or CML group (Fig. 4b-e), which further demonstrated the crucial role of the oral delivery system in facilitating the efficacy of *EcN*. Furthermore, RT-PCR was used to reveal the molecular mechanism of CML@*EcN* on repairing gut barrier. CML@*EcN* Significantly upregulated the expression levels of ZO-1 and Occludin in the colon tissue of UC mice (Fig. 4f-h). H&E staining was employed for further investigate the repair effect of CML@*EcN* on the epithelial barrier of colon (Fig. 4i). Horizontally stained images revealed the overall histological characteristics of the colon tissues in each group of mice. Compared to the Healthy group, UC-bearing mice without treatment showed separation between the colon epithelium and surrounding tissues (highlighted by black arrows), with the colon epithelium appearing irregular. CML@*EcN* significantly improved these pathological features, with the colon epithelium closely connected to surrounding tissues, and the brush border structure of the colon epithelium appearing clear and intact (delineated by dashed lines). Besides, the localized magnified horizontal and longitudinal sections showed that the colon tissue of Healthy group mice is arranged in a regular and dense U-shape, and the crypt and colon epithelial barrier were clearly visible (highlighted by red arrows). However, the induction of DSS made the contour of colon epithelial tissue completely indistinguishable, and the whole colon tissue was diffuse, which indicated that the colon tissue of UC-bearing mice had been seriously damaged. Compared to the other three intervention groups and positive drug groups, CML@*EcN* best maintained the integrity of colon epithelial tissue in mice. Furthermore, the MPO staining results indicated that, the damage to the gut barrier in UC mice led to obvious infiltration of neutrophils (highlighted by red dashed lines, Fig. 4j), while CML@*EcN* significantly eliminated this phenotype.

**Fig. 4.**
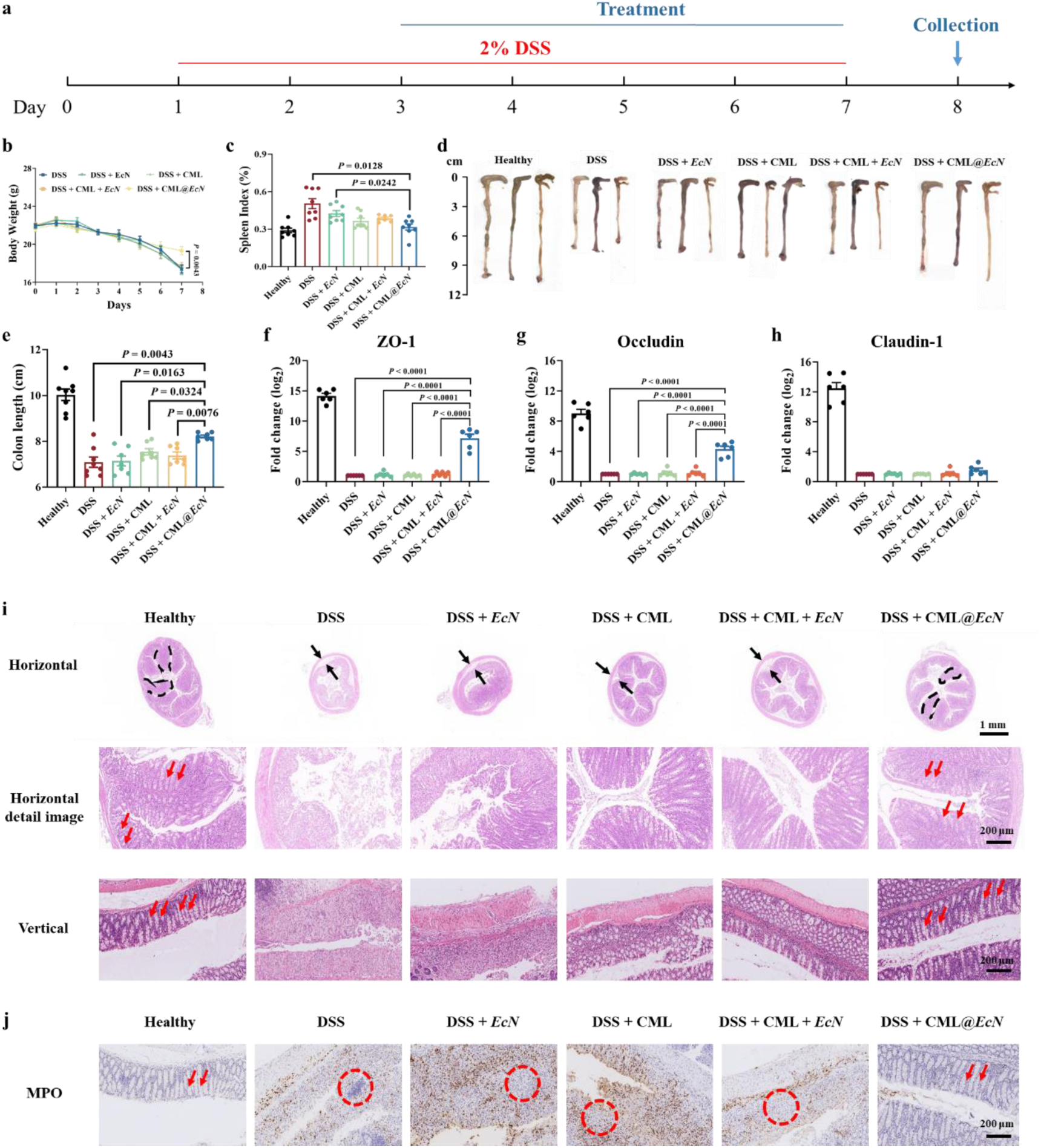
Evaluation of CML@EcN for UC treatment and gut barrier repair efficacy. **a,** Schematic of *in vivo* experimental. After 7 days of modeling and intervention, CML@*EcN* significantly alleviated the symptoms of **b,** weight loss, **c,** increased spleen weight index and **d,e,** shortened colon length in DSS model, free *EcN* and CML group. **f-h,** CML@*EcN* significantly increased the level of ZO-1 and occludin in colon tissue (n = 6 independent animals in **b, c, e – h,** n = 3 independent animals in **d**). **i,** Colons of each group of mice were sliced horizontally in overall image. Scale bar, 1 mm, and longitudinally histological observations were performed using HE staining (The brush border of colon is highlighted by dashed areas or red arrows). Scale bar, 200 μm. **j,** MPO staining showed that CML@*EcN* significantly repaired the colon epithelial barrier and avoided the infiltration of neutrophils in the colon tissue (The area of neutrophil infiltration is highlighted by red arrows). Scale bar, 200 μm.

All the results suggested that CML@*EcN* significantly repaired the gut barrier in UC mice by regulating the expression of ZO-1 and occludin in the colon tissues. Normal expressions of ZO-1 and occludin are directly related to the proliferation of intestinal epithelium and the integrity of the gut barrier^16, 17^, revealing that CML@*EcN* can obviously repair the intestinal barrier at the molecular level.

### CML@*EcN* can modulate the immune homeostasis of UC-bearing mice

UC-induced damage of gut barrier also leads to a large amount of GM entering the surrounding tissues, triggers an immune response and causes excessive immune activation in the local intestinal environment, followed by, resulting in a large infiltration of immune cells in the gut lumen, exacerbating the disruption of the gut environment, forming severe immune disorder^18, 19^. In present study, CML@*EcN* had significant regulatory effects on peripheral blood, colon, and spleen immunity in UC mice. In the peripheral blood, UC mice showed a significant increase in the pro-inflammatory cytokines IL-6, TNF-*α*, and IL-1*β* concentrations, along with a significant decrease in the anti-inflammatory cytokine IL-10 concentration (Fig. 5a-d, Supplementary Fig. 15). CML@*EcN* significantly downregulated the concentration of IL-6 and dramatically upregulated the concentration of IL-10 (Fig. 5a-d). Furthermore, single-cell flow cytometry was used to classify immune cells in the colon and spleen tissues of mice in each group. The results indicated that severe immune dysregulation was observed in the colon and spleen of UC mice, with a severe imbalance in the proportion of neutrophils, T cells, B cells, macrophages, and dendritic cells in the colon (Fig. 5e-i, Supplementary Fig. 16-20). CML@*EcN* significantly downregulated the content of neutrophils (9.0% vs 11.0%, *P* = 0.0079) and B cells (17.6% vs 31.1%, *P* = 0.0266), and significantly upregulated the T cell content (61.7% vs 60.1%, *P* = 0.011) in the colon of UC mice. Furthermore, aside from the colon, the spleen plays a crucial role in the immune regulation of UC. The migration of T cells from the spleen to the colon is closely associated with gut barrier repair^20, 21^. In present study, the proportion of CD3^+^CD4^+^ and CD3^+^CD8^+^ T cells significantly decreased in UC mice. CML@*EcN* significantly upregulated the proportion of CD3^+^CD4^+^ (25.2% vs 15.3%, *P* = 0.0416) and CD3^+^ CD8^+^ (39.7% vs 18.8%, *P* = 0.0055) T cells, while the other intervention groups did not show similar effects (Fig. 5j-m).

**Fig. 5.**
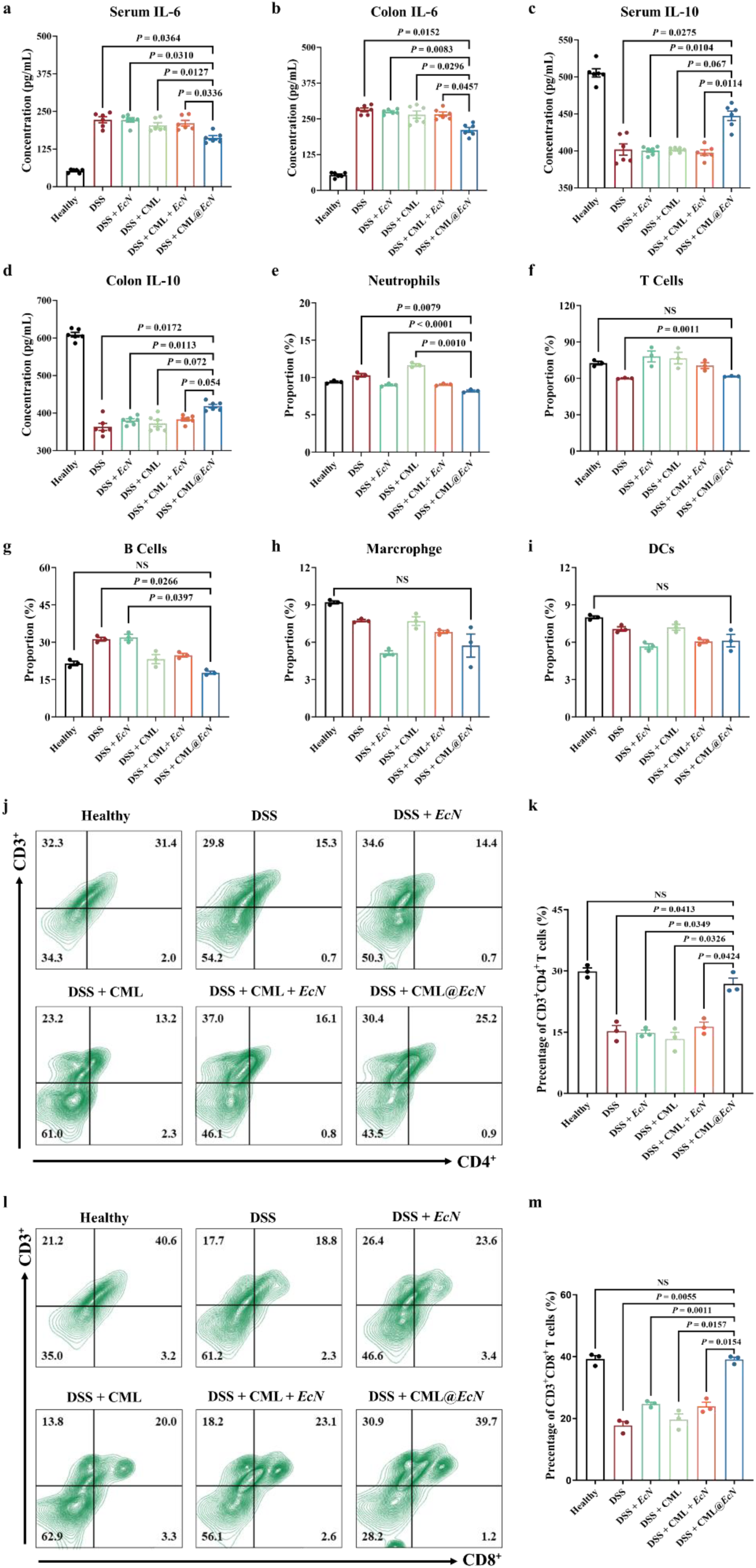
CML@*EcN* effectively regulated the immune homeostasis *in vivo*. After 7 days of intervention, **a-d,** CML@*EcN* Significantly improved the levels of inflammatory factors in peripheral blood and colon tissue. **e-i,** single-cell flow cytometry of colon tissue was used to perform immune cell typing in colon tissue of mice in each group. Indicating CML@*EcN* significantly improved the content of neutrophils, T cells, and B cells in the colon of UC mice. Furthermore, flow cytometry was used to classify immune cells in the spleen of mice in each group, and CML@*EcN* significantly upregulated the proportion of **j,l,** CD3+CD4+and **k,m,** CD3+CD8+ T cells in the spleen of UC mice.

The results provide strong evidence that CML@*EcN* effectively modulates the immune system in UC mice by regulating peripheral blood inflammatory factors and the composition of immune cells in the colon and spleen. Specifically, CML@*EcN* enhances the circulation of T cells between the spleen and gut, suppresses the abnormal immune microenvironment activated in the colon, promotes intestinal barrier repair, reduces the ratio of neutrophils and B cells in the colon, and restores immune balance between the colon and spleen during UC treatment.

### CML@*EcN* reshaped the gut environment of UC mice

In addition to the damage to the epithelial barrier of the colon and the body’s immune system, UC can also cause serious gut environment (GM & metabolites structural) disorders^22, 23^. Therefore, 16S rRNA and metabolomics sequencing were used respectively to evaluate the impact effect of CML@*EcN* on GM and metabolites structure in DSS model mice.

In terms of GM structure, after 7 days of modeling and intervention, the DSS model mice showed severe GM disorder, while CML@*EcN* reshaped the GM structure Specifically, the GM α-diversity of DSS mice was significantly improved by CML@*EcN* (Fig. 6a). In the overall structure, the GM structure of DSS mice was seriously disordered, forming a new group different from normal mice, while the intervention of CML@*EcN* made the GM structure of mice similar to that of normal mice (Fig. 6b, Supplementary Fig. 21b). The differential flora of GM in each group of mice was further excavated. At the phylum level, the percentage stacking histogram visually showed that CML@*EcN* has basically recovered the GM structural disorder caused by DSS (Supplementary Fig. 21a). Importantly, Firmicutes and Bacteroidetes occupy a large proportion in the whole GM, and the changes in their abundance usually reflect the status of body’s inflammation, immune and metabolic system. In this work, CML@*EcN* effectively decreased the abundance of Firmicutes and increased the abundance of Bacteroidetes in the faeces of mice (Supplementary Fig. 21c,d). At the genus level, further validation was conducted using FISH to assess the content of gut microbiota (GM) related to gut barrier repair and immune regulation (Fig. 6d, Supplementary Fig. 22, 23). The results indicated that CML@*EcN* significantly upregulated the abundance of *Akkermansia* (11.82% *vs* 4.12%), *Enterocloster* (3.47% *vs* 0.75%), *Turisimonas* (1.61% *vs* 0.46%), and *Muribaculum* (7.46% *vs* 1.83%) in the gut of UC mice (Fig. 6e). Correspondingly, CML@*EcN* also reshaped the structure of metabolites in the colon contents of DSS mice, with over 400 metabolites significantly upregulated and over 100 metabolites significantly downregulated (Fig. 6f,g).

**Fig. 6.**
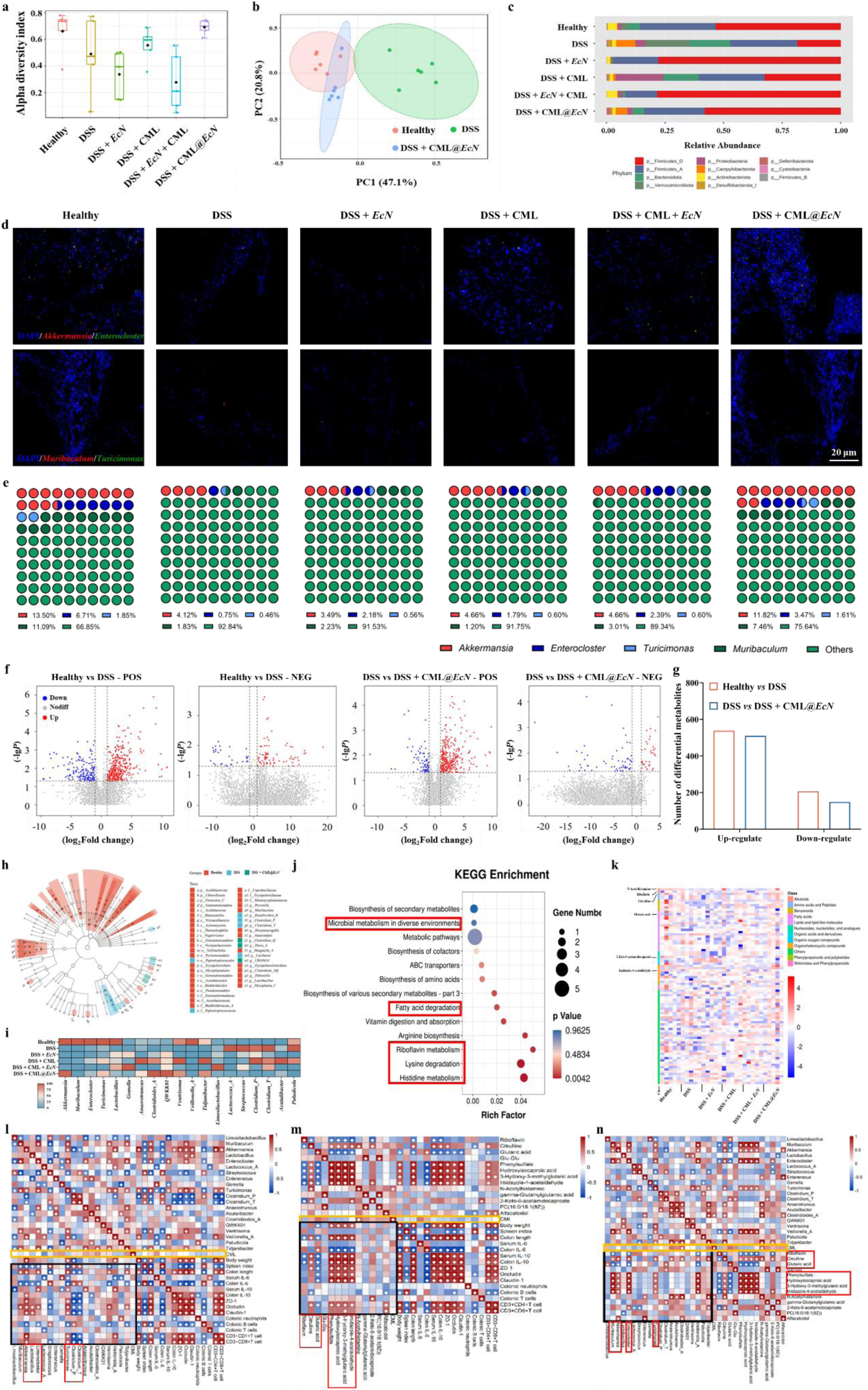
CML@*EcN* reshaped the gut environment of UC mice and modulated immune homeostasis *via* metabolism. **a,b,** After 7 days of intervention, CML@*EcN* Significantly improved the Alpha and Beta diversity of GM in UC mice. **c,** The percentage stacking histogram visually showed the phylum GM composition of mice in each group. **d,** Double labeled FISH showed *Akkermansia* & *Enterocloster*, *Muribaculum* & *Turisimonas* in each group, respectively. Scale bar, 20 μm. **e,** Quantitative analysis of FISH. **f,g,** CML@*EcN* Improved the metabolite structure of colon contents. **h,** LDA analysis was used to further screen the key differential flora at the genus level. **i,** Heat map of differential GM. **j,** Functional annotation of different metabolites and key pathways. **k,** the differential metabolites involved in these key pathways are highlighted in the heat map. **l-n,** correlation analysis was used to analyze the relationships between GM & metabolites, GM & efficacy/mechanisms, and metabolite & efficacy/mechanisms, respectively.

To further identify the key microbial communities and metabolites involved in repairing the intestinal barrier and maintaining immune homeostasis, Linear Discriminant Analysis Effect Size (LEfSe) was employed to recognize significantly differentially abundant GM groups among the categories and to assess the contribution of these differences to sample classification (Fig. 6h). The differential flora with LDA > 3, and *P* < 0.05 was extracted. Furthermore, the differential flora that CML@*EcN* reversing the change trend of DSS was screened (Fig. 6h,i). It should be noted that sequencing results did not show a significant increase in the abundance of *EcN*, which may have limited by the sequencing method used in this study. 16S rRNA sequencing could not accurately identify all microorganisms at the level of species or even strains, so *EcN* information could not be detected^24^. On the other hand, FISH analysis detected *EcN* in the colonic contents of mice (Fig. 3c,d), which served as conclusive evidence of successful colonization of *EcN* in the mouse colon, further corroborated by the results from IVIS imaging (Fig. 3a,b).

Furthermore, match these key differential metabolites with the HMDB database for type annotation, and with the KEGG database for functional annotation. The results indicated that these differential metabolites are closely associated with the metabolism of fatty acids, amino acids, and other substances in the body (Fig. 6j). Furthermore, six important differential metabolites (N-acetylhistamine, riboflavin, citrulline, glutaric acid, 2-keto-6-acetamidocaproate, imidazole-4-acetaldehyde) involved in the aforementioned life processes were ultimately screened out (Fig. 6k).

Among the GMs and metabolites we have selected, *Muribaculum* is associated with host inflammation and intestinal immune response, and an increase in *Muribaculum* abundance contributes to the alleviation of UC symptoms^25^; *Enterocloster* can induce gut epithelial immune responses and play an important role in combating pathogenic bacterial infections^26^; As a widely recognized probiotic, *Akkermansia* has been reported to alleviate UC symptoms, and its efficacy in restoring intestinal barrier and improving immune system function has been confirmed^27, 28^. Furthermore, key differential metabolites primarily associated with host vitamin, fatty acid, and amino acid metabolism pathways (Fig. 6j), which suggested that CML@*EcN* may achieve its therapeutic effects in treating UC by regulating nutrient metabolism.

### Correlation among GM, metabolites, efficacy and mechanisms of UC treatment

To correlate CML@*EcN*, UC phenotype, immune system function and gut environment, correlation analysis was used to reveal the internal relationship among them (Fig. 6i-n). GM and metabolites which related to inflammation, immune system and colon barrier repair (*Muribaculum*, *Enterocloster*, *Akkermansia*, *Turicimonas* and riboflavin, glutaric acid, citrulline and imidazole-4-acetaldehyde) were strong positive correlation with body weight, colon length, serum and colon tissue anti-inflammatory factor IL-10 level, expression levels of ZO-1 and Occludin in colon tissue and immune regulation in colon and spleen and strong negative correlation with pro-inflammatory factor IL-6 level (The black and red boxes in Fig. 6i-n). Additionally, CML did not show a strong correlation with any factors (The yellow boxes in Fig. 6i-n), which indicated that in CML@*EcN*, the therapeutic effects for UC solely originate from *EcN*. CML plays a role in protecting *EcN* and facilitating its pH responsive precise release in the colon. In fact, CML is highly resistant to degradation in the mammal body^29, 30^, in present study, over 70% of CML was excreted with the faeces (Supplementary Fig. 14), indicating that CML does not accumulate in the body or participate in organism metabolism, posing no biosafety hazards.

Integration through correlation analysis, several clues of "*EcN*-GM-metabolites-efficacy/mechanism" have been extracted. In terms of vitamin metabolism: *EcN* upregulates the abundance of *Muribaculum/Intercloster,* and subsequently downregulates the abundance of riboflavin, achieving the effect of regulating peripheral blood immunity; in terms of fatty acid metabolism: *EcN* upregulates the abundance of *Muribaculum/Intercloster,* and subsequently downregulates the abundance of glutaric acid, achieving the effect of regulating peripheral blood immunity. Besides, *EcN* upregulates the abundance of *Akkermansia* and subsequently downregulates the abundance of glutitic acid, achieving the function of repairing the colon barrier/regulated B and T cells; in terms of amino acid metabolism: *EcN* upregulates the abundance of *Muribaculum/Intercloster*, thereby increasing the abundance of citrulline and imidazole-4-acetaldehyde, achieving the effect of regulating immunity. Besides, *EcN* upregulates the abundance of *Akkermansia/Turicimonas*, thereby increasing the abundance of citrulline and imidazole-4-acetaldehyde, achieving the effect of repairing the colon barrier/regular immunity.

### Outlook

In present study, we designed and fabricated an orally surface engineered probiotics delivery system CML*@EcN* as ballistic helmet, which maintained the high activity of *EcN* in the gastrointestinal tract while achieved the release profile on-demand and efficient colonization of *EcN* at the site of colitis. Interestingly, the release and proliferation process of *EcN* in the colon can be modeled as a plug-flow mode We derived a mathematical expression for the release and proliferation of *EcN* over time using the plug-flow model, which provides a theoretical basis for the design and preparation of similar oral probiotic formulations. Furthermore, CML@*EcN* with orally and daily administrated lower doses showed satisfactory UC treatment results (much lower than another commercially available oral probiotic preparation for UC treatment (VSL#3, oral dose 10^11^ CFU)^31, 32^). In terms of mechanism research, we revealed the therapeutic mechanism of CML@*EcN* in immune modulation and gut environment reshaping, and linked *EcN*, GM, metabolites, efficacy and mechanisms from the perspectives of vitamin, fatty acid, and amino acid metabolism. In other words, we successfully explored the relationship between immune homeostasis and GM reconstitution via metabolism, which opened this black box. In future, we would build on the models and mechanisms revealed in present study to expand the variety of probiotics, and develop a series of colon-targeted oral probiotic formulations for different diseases treatment.

In summary, the dynamic model and formulas constructed in this work are universality and provide reference for the rational design of similarly precise colon controlled release oral probiotic delivery systems. The therapeutic mechanism of CML@*EcN* revealed in this work provides inspiration for the development of new targets and therapies for UC treatment, as well as new directions for the treatment of colon diseases by orally administered probiotics *via* modulating GM and immune homeostasis.

## Methods

### Materials

*Escherichia coli* Nissle 1917 was purchased from Mingzhou Biotechnology Co., Ltd., mCherry labeled *EcN* was purchased from Beijing Zhuangmeng International Biogene Technology Co., Ltd, lux-reporter plasmid *EcN* was constructed by Hangzhou Baosai Biotechnology Co., Ltd, tryptone and sodium chloroacetate were purchased from Shanghai Aladdin Biochemical Technology Co., Ltd., sodium chloride, yeast extract, and folinol reagent were purchased from Shanghai Macklin Biochemical Co., Ltd. Alkali lignin was purchased from Zhejiang Jiefa Technology Co., Ltd.

### Modification and validation of lignin

#### Modification of lignin

Refer to the method of Chen et al. and make slight improvement to modify lignin^33^. Briefly, 7.05g alkali lignin is dissolved in 28mL NaOH solution (pH=12), and ClCH_2_COONa aqueous solution (25wt.%) is dripped with peristaltic pump under stirring. The reaction is carried out in an oil bath at 85 ℃, and the pH value of the reaction solution is always controlled to be greater than 11. After the dripping of ClCH_2_COONa aqueous solution, the reaction continues at 85 ℃ for 3h. After the reaction, the reaction solution is dialyzed for one week, and then concentrated and dried to obtain carboxymethyl alkali lignin (CAL). Change the amount of ClCH_2_COONa added to obtain CAL1, CAL2 and CAL3, as shown in Supplementary Table 1.

### Fourier transform infrared spectroscopy (FTIR)

The infrared spectra of AL, CAL1, CAL2 and CAL3 were measured with the Tensor 27 Fourier infrared spectrometer (Bruker, Germany). The scanning wavenumber range was 4000-400cm-1, and the resolution was 4cm-1. The selected test method is potassium bromide tablet method.

### Aqueous phase potentiometric titration

The carboxyl content of four kinds of lignin was tested with the 809 Titrando automatic potentiometric titrator (Mertrohm, Switzerland), and the titrant used was HCl standard solution. 30mg of lignin is dissolved in 5mL of KOH solution (pH=14), then 50mg of p-hydroxybenzoic acid and 25mL of deionized water are added, and the test is started after 30min of ultrasound. This is the experimental group. Other conditions remain the same, and the solution without lignin is used as the blank group. The calculation formula of carboxyl content is as follows^34^:

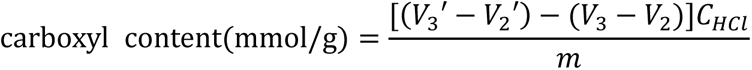

In the formula, V_2_’, V_3_’, V_2_ and V_3_ are the volume of titrant consumed at the second and third break points on the titration curve of the experimental group and the blank group, mL; C_HCl_ is the concentration of HCl standard solution, mmol/L; m is the absolute dry mass of lignin sample, g.

### Folin−Ciocalteu’s phenol (FC) method

The phenolic hydroxyl content of lignin was measured by UV-visible spectrophotometer (Shimadzu UV-2450, Japan). Dissolve the dried lignin in the alkaline aqueous solution, then add the Folin−Ciocalteu’s phenol reagent, and then add the Na_2_CO_3_ solution (20 wt.%) after a period of time. Test the absorption value of the mixed solution at 760 nm. Substitute the absorption value into the standard equation to calculate the phenol hydroxyl content^35^.

### Fabrication of *EcN* oral delivery system

According to the principle of electrostatic adsorption, firstly, add 1 mL of CaCl_2_ solution to 1 mL of bacterial solution, incubate for 15 min, centrifuge at 1000 rpm for 6 min, remove the supernatant, and then wash twice with sterile water to fully remove the unabsorbed CaCl_2_. Then add 1mL of carboxymethyl alkali lignin solution to the precipitate obtained after centrifugation, incubate for 15min, and centrifuge at 10000rpm for 6 minutes to remove the unabsorbed carboxymethyl alkali lignin, wash it with sterile water twice, finally re-suspend the bacteria in phosphate buffer salt solution (PBS), and store it in a refrigerator at 4 ℃ for subsequent dynamic light scattering (DLS) (ZS Nano S, Brookhaven, USA) test.

### Scanning electron microscope (SEM) and transmission electron microscopy (TEM) observation

The surface morphology of bacteria was observed by field emission electron SEM (HITACHI SU8220, Japan). Centrifuge the bacterial suspension at 10000 rpm for 3min, remove the supernatant, add 2.5wt.% glutaraldehyde solution, soak overnight to fix the bacterial form, then remove the glutaraldehyde solution by centrifugation, use 10%, 30%, 50%, 70%, 90% ethanol aqueous solution for gradient elution treatment, treat each gradient for 1h, then remove the supernatant by centrifugation, add the next concentration gradient of ethanol solution for further treatment, and finally suspend the bacteria in 100% ethanol solution, Take 50 µL bacterial suspension and dry it in a vacuum oven on a monocrystalline silicon wafer, and then use a scanning electron microscope to photograph the surface morphology of the sample. For TEM (FEI spirit T12), the sample suspension was dropped onto a copper grid and tested after air-drying overnight.

### Laser scanning confocal microscopy image

Confocal microscopy was utilized for imaging of staining of live and dead bacteria (live & dead bacterial staining kit, Yisheng Biotechnology (Shanghai) Co., Ltd) as well as CML@ *EcN* (mCherry-labeled). After sample preparation, fixation was performed using 4% PFA. The fixed samples were then dropped onto glass slides, and sealing, observation and photography were conducted using confocal microscopy (Nikon, AIR).

### Simulated gastric environment experiment

Prepare simulated gastric juice (0.15 M HCl, 0.05 M KCl, pH=1.2) according to the relevant reported methods^36^. Take the wrapped and unwrapped *E Coli* Nissle 1917 (OD=0.4) 5 ml, add 10 ml of simulated gastric juice and PBS (pH=7.4) respectively, stir with magnetic force, take 1ml of bacterial solution at 30 min, 60 min, 90 min and 120 min respectively, centrifuge at 4200 rpm for 10 min, wash with LB medium once, re-suspend with LB medium and transfer into 15 ml centrifuge tube, incubate with shaking bed at 37 ℃ for 24h, and measure the OD_600_ value of each sample respectively.

### *In vivo* biological distribution experiment

Lux-reporter plasmid *EcN* was utilized to prepare CML@*EcN*, which was then administered *via* gavage to mice. At 4 hours and 24 hours’ post-gavage, *in vivo* imaging system (IVIS, BrukerIn-Vivo F PRO) was performed on the mice, and the intestines and vital organs were extracted for IVIS imaging. Quantitative analysis of bioluminescent signals was conducted using the software’s built-in modules.

### Biosafety experiment

Human normal colon epithelial cells (NCM-460) were used as a cell model for safety evaluation *in vitro*^37^. NCM-460 cells were maintained in DMEM supplemented with 10% (v/v) FBS and anti-biotics (1% penicillin and 1% streptomycin) at 37 °C in a humidified atmosphere containing 5% CO_2_. These cells were seeded in pre-adipocyte medium into 96-well tissue culture plates at approximately 8 × 10^3^ cells/well and grown to confluency; after confluency, incubate different concentration delivery systems without *EcN* with cells for 24 h and 48 h respectively. Cell Counting Kit-8 (CCK-8, Sangon Biotech, Shanghai, China) is used to detect the cytotoxicity.

Male C57BL/6N mice were randomly divided into three groups based on gavage material: the PBS group, the CML low-dose group (800 mg/L), and the CML high-dose group (2400 mg/L), with n=6 in each group. They were gavaged continuously for 2 weeks. On the 15th day of the experiment, blood samples were collected from each group of mice for biochemical analysis. Important organs (intestine, heart, liver, spleen, lung, and kidney) were collected and sliced for H&E staining to observe tissue morphology.

### *In vivo* efficacy and mechanism verification

#### Animal care and *in vivo* experiment procedures

All experimental procedures were conducted and the animals were used according to the Guide for the Care and Use of Laboratory Animals published by the institutional animal care and use committee (IACUC) and approved by the Animal ethics committee of Tsinghua University (Approve ID: 2023F133). Male C57BL/6N mice (6 weeks old) were purchased from Vital River Laboratories (Beijing, China) and housed (three animals/cage) under 22 ± 2 ℃ and 55% ± 10% humidity with 12 h light/12 h dark cycle in a specific pathogen-free (SPF) animal room. Animals were acclimated to the environment for 7 days with normal commercial basic diet and sterile water ad libitum before the experiment.

As shown in Fig. 5A. Six mice were randomly selected to maintain normal drinking water as the control group (Control, n = 6), and the other mice were induced UC with 2% DSS. intervened free *EcN* (DSS + *EcN*), CML (DSS + CML), *EcN* and CML mixed directly (DSS + CML + *EcN*), and CML@*EcN* (DSS + CML@*EcN*). The experiment lasted on the 15th day, and mice were euthanized, colon, spleen and blood of mice were collected for further analysis.

### Hematoxylin and eosin & myeloperoxidase (MPO) staining

The method of H & E staining refers to exist reports^38, 39^. Briefly, the tissue was fixed in 4% paraformaldehyde for paraffin section. Firstly, the slices were dyed in hematoxylin solution for a few min and separated in acid water and ammonia water for a few seconds. After that, the slices were washed with running water for 1 h, dehydrated in 70% and 90% ethanol for 10 min, and then stained in eosin staining solution for 2 to 3 min. The stained sections were dehydrated with pure ethanol and then penetrated with xylene. Finally, the transparent section was dropped with gum and sealed with cover glass.

The method of myeloperoxidase staining refers to the relevant process of immunohistochemistry^40^. Briefly, after the tissue section is dewaxed, the antigen is repaired with citric acid (pH = 6), the antibody is sealed and incubated, and then it is re-stained with hematoxylin, dehydrated and dried with gradient alcohol, and photographed after the tablet is sealed.

### Inflammatory factor detection

The colon tissues were ground with a tissue grinder for 1min, the tissue homogenate was taken for ELISA detection, and the mouse blood was left at room temperature for 2-4 h, centrifuged at 3500 rpm for 15 min, and the upper serum was taken for subsequent detection. The concentration of inflammatory factors was detected with the corresponding ELISA kit (Thermo Fisher Scientific, Massachusetts, USA).

### Colonic immune cell single-cell flow cytometry

Transfer the intestinal tissue to a 50 mL centrifuge tube containing 10 mL of separation solution, and place it in a constant temperature shaking incubator at 37°C with agitation at 250 rpm for 15 minutes. During this period, vigorously shake the tube once to remove intestinal epithelial cells. If the separation solution becomes turbid, proceed to the next step; if the solution remains clear, continue shaking for an additional 5-10 minutes until the solution becomes turbid. Stop shaking when the separation solution becomes turbid. Wash the remaining tissue with 10 mL of D-Hanks solution (Wuhan Pricella Biotechnology Co., Ltd.) to remove residual separation solution. Blot dry on paper, then cut into a paste with scissors and transfer to a 50 mL centrifuge tube. Add 5 mL of dissociation solution and shake at 37°C with agitation at 250 rpm for 30 minutes. Subsequently, filter through a 70 μm filter membrane at 4°C and 500g for 15 minutes. Finally, resuspend cells in 3-5 mL of FACS buffer (Beijing Tianjingsha Gene Technology Co., Ltd), vortex to mix thoroughly, and incubate with appropriate antibodies for flow cytometric analysis (antibody information provided in Supplementary Table 3).

### Spleen immune cell single-cell flow cytometry

The spleen was minced and passed through a 70-micron filter at 4°C and 400g for 10 minutes. After discarding the supernatant, red blood cell lysis buffer was added, followed by incubation on ice for 15 minutes. Subsequently, staining with CD4 and CD8 antibodies (Supplementary Table 3), followed by three washes with PBS. Cells were then resuspended in flow cytometry buffer for test.

### Quantitative real-time reverse-transcription PCR

Tissue RNA was extracted by Trizol method. Briefly, the tissue was collected in a 2ml RNase free centrifuge tube, then, Trizol was added to centrifuge, tissue grinding fluid was obtained by tissue grinder (Hede Technology, Beijing. N9548), centrifuged (12000 rpm, 5 min) to take the supernatant, added 1 mL chloroform, fully mixed, placed at room temperature for 15min, centrifuged to take the supernatant (12000 rpm, 15 min), added 600 μL isopropanol, placed on ice for 20min, centrifuged to discard the supernatant (12000 rpm, 20 min), precipitated was washed with ethanol, and dried in the fume hood, finally, add 50 μL DEPC water and store at - 80 ℃.

Equal amounts of RNA to synthesize cDNA with the transScript One-Step gDNA Removal and cDNA Synthesis SuperMix kit (AT311-03, TransGen Biotech, Beijing, China). Quantitative real-time PCR (qRT–PCR) was performed in triplicate using SYBR Green, 96-well plates and the Real-Time PCR System (Bio-Rad, Hercules, CA, USA). Each well was loaded with a total of 20 μL containing 2 μL of cDNA, 2 x 0.4 μL of target primers, 7.2 μL of water and 10 μL of SYBR Fast Master Mix. We performed hot-start PCR for 45 cycles, with each cycle consisting of denaturation for 5 s at 95 ℃, annealing for 15 s at 58 ℃ and elongation for 10 s at 72 ℃. GAPDH expression was used to normalize the mRNA expression. The primers used in present research are shown in Supplementary Table 4.

### 16S rRNA sequencing analysis

At the end of the experiment, the fresh mouse faeces were collected into a sterile centrifuge tube with sterile tweezers and stored at - 80 ℃. Then, fecal samples were send to Parsenor Biotechnology Co., Ltd. with dry ice for 16S rRNA sequencing.

### Extraction of genome DNA

Total genome DNA from samples was extracted according to manufacturer’s protocols. DNA concentration was monitored by Equalbit dsDNA HS Assay Kit.

### Amplicon Generation & Library preparation

20-30 ng DNA was used to generate amplicons. V3 and V4 hypervariable regions of prokaryotic 16S rDNA were selected for generating amplicons and following taxonomy analysis. GENEWIZ designed a panel of proprietary primers aimed at relatively conserved regions bordering the V3 and V4 hypervariable regions of bacteria and Archaea16S rDNA. Then, a linker with Index is added to the end of the PCR product of 16S rDNA by PCR for NGS sequencing, the library is purified with magnetic beads, and the concentration is detected by a microplate reader and the fragment size is detected by agarose gel electrophoresis.

### Illumina sequencing

Detect the library concentration by a microplate reader. The library was quantified to 10nM, and PE250/FE300 paired-end sequencing was performed according to the Illumina MiSeq/Novaseq (Illumina, San Diego, CA, USA) instrument manual. The MiSeq Control Software (MCS)/Novaseq Control Software (NCS) Read sequence information.

### Data analysis

The raw data were uploaded to the MicrobiomeAnalyst platform (https://www.microbiomeanalyst.ca/) for GM structure, differential flora analysis and functional annotation of mice in each group.

### Metabolomics sequencing analysis

Colonic contents samples of each groups mice were randomly selected and s send to Parsenor Biotechnology Co., Ltd. with dry ice for. metabolomics analysis. A 400 μL solution (methanol: water =7:3, V/V) containing internal standard was added into 20 mg sample, and vortexed for 3 min. The sample was sonicated in an ice bath for 10 min and vortexed for 1 min, and then placed in −20 ℃ for 30 min. The sample was then centrifuged at 12000 rpm for 10 min (4 ℃). And the sediment was removed, then centrifuged the supernatant at 12000 rpm for 3 min (4 ℃). A 200 μL aliquots of supernatant were transferred for LC-MS analysis. All samples were acquired by the LC-MS system followed machine orders. The analytical conditions were as follows, UPLC: column, Waters ACQUITY UPLC HSS T3 C18 (1.8 μm, 2.1 mm*100 mm); column temperature, 40℃; flow rate, 0.4 mL/min; injection volume, 2 μL; solvent system, water (0.1% formic acid): acetonitrile (0.1% formic acid); gradient program, 95:5 V/V at 0 min, 10:90 V/V at 11.0 min, 10:90 V/V at 12.0 min, 95:5 V/V at 12.1 min, 95:5 V/V at 14.0 min. Data analysis and mapping through were completed on the Paisenor Gene Cloud platform (https://www.genescloud.cn/home).

### Statistical analysis

For statistical differences analysis between groups, two way ANVOA was used to analyze the weight data of mice, and other data were analyzed with one way ANVOA. Data were presented as mean ± SEM. Significant differences were considered when P < 0.05. Graph-Pad Prism 9 (GraphPad Software, San Diego, CA, USA) was used for data analysis.

## Acknowledgements

This work was supported by the Key Research and Development Program of the Ministry of Science and Technology [2023YFA0913600 (2023YFA0913602)], the Shenzhen Medical Research Fund (SMRF: B2302009), the National Natural Science Foundation of China (22278242), the Funds from Shenzhen International Graduate School at Tsinghua University (HW2023009, JC2021011).

## Reference

1. Peterson, L.W. & Artis, D. Intestinal epithelial cells: regulators of barrier function and immune homeostasis. Nature reviews immunology 14, 141–153 (2014).

2. Kudelka, M.R., Stowell, S.R., Cummings, R.D. & Neish, A.S. Intestinal epithelial glycosylation in homeostasis and gut microbiota interactions in IBD. Nature reviews Gastroenterology & hepatology 17, 597–617 (2020).

3. Nowarski, R. et al. Epithelial IL-18 equilibrium controls barrier function in colitis. Cell 163, 1444–1456 (2015).

4. Mehandru, S., Allen, P., Peyrin-Biroulet, L. & Colombel, J. Ulcerative colitis. The Lancet 10080, 1756–1770 (2017).

5. Fugger, L., Jensen, L.T. & Rossjohn, J. Challenges, progress, and prospects of developing therapies to treat autoimmune diseases. Cell 181, 63–80 (2020).

6. Zhou, J. et al. Programmable probiotics modulate inflammation and gut microbiota for inflammatory bowel disease treatment after effective oral delivery. Nature Communications 13, 3432–3455 (2022).

7. Sanders, M.E., Merenstein, D.J., Reid, G., Gibson, G.R. & Rastall, R.A. Probiotics and prebiotics in intestinal health and disease: from biology to the clinic. Nature reviews Gastroenterology & hepatology 16, 605–616 (2019).

8. Losurdo, G., et al. *Escherichia coli* Nissle 1917 in ulcerative colitis treatment: systematic review and meta-analysis. J Gastrointestin Liver Dis 24, 499–505 (2015).

9. Heavey, M.K. et al. Targeted delivery of the probiotic Saccharomyces boulardii to the extracellular matrix enhances gut residence time and recovery in murine colitis. Nature Communications 15, 3784–3799 (2024).

10. Liu, J. et al. Gut lumen-targeted oral delivery system for bioactive agents to regulate gut microbiome. Journal of Future Foods 2, 307–325 (2022).

11. Wang, X. et al. Bioinspired oral delivery of gut microbiota by self-coating with biofilms. Science Advances 6, eabb1952 (2020).

12. Cao, Z., Wang, X., Pang, Y., Cheng, S. & Liu, J. Biointerfacial self-assembly generates lipid membrane coated bacteria for enhanced oral delivery and treatment. Nature Communications 10, 5783–5793 (2019).

13. Chen, Y. et al. Reinforcement of the intestinal mucosal barrier via mucus-penetrating PEGylated bacteria. Nature Biomedical Engineering 12, 1–19 (2024).

14. Kirby, R. Actinomycetes and lignin degradation. Advances in applied microbiology 58, 125–168 (2005).

15. Kaps, M. & Lamberson, W.R. Biostatistics for animal science. (CABI, 2004).

16. Kuo, W.T., Odenwald, M.A., Turner, J.R. & Zuo, L. Tight junction proteins occludin and ZO-1 as regulators of epithelial proliferation and survival. Annals of the New York Academy of Sciences 1514, 21–33 (2022).

17. Otani, T. & Furuse, M. Tight junction structure and function revisited. Trends in cell biology 30, 805–817 (2020).

18. Zundler, S. et al. Gut immune cell trafficking: inter-organ communication and immune-mediated inflammation. Nature Reviews Gastroenterology & Hepatology 20, 50–64 (2023).

19. Tian, Z. et al. Gut microbiome dysbiosis contributes to abdominal aortic aneurysm by promoting neutrophil extracellular trap formation. Cell host & microbe 30, 1450–1463 (2022).

20. Sharma, A., Tirpude, N.V., Kulurkar, P.M., Sharma, R. & Padwad, Y. Berberis lycium fruit extract attenuates oxi-inflammatory stress and promotes mucosal healing by mitigating NF-κB/c-Jun/MAPKs signalling and augmenting splenic Treg proliferation in a murine model of dextran sulphate sodium-induced ulcerative colitis. European journal of nutrition 59, 2663–2681 (2020).

21. Tilg, H., Adolph, T.E. & Trauner, M. Gut-liver axis: Pathophysiological concepts and clinical implications. Cell Metabolism 34, 1700–1718 (2022).

22. Lavelle, A. & Sokol, H. Gut microbiota-derived metabolites as key actors in inflammatory bowel disease. Nature reviews Gastroenterology & hepatology 17, 223–237 (2020).

23. Schirmer, M. et al. Compositional and temporal changes in the gut microbiome of pediatric ulcerative colitis patients are linked to disease course. Cell host & microbe 24, 600–610 (2018).

24. Muhamad Rizal, N.S., et al. Advantages and limitations of 16S rRNA next-generation sequencing for pathogen identification in the diagnostic microbiology laboratory: perspectives from a middle-income country. Diagnostics 10, 816 (2020).

25. Shaler, C.R. et al. Psychological stress impairs IL22-driven protective gut mucosal immunity against colonising pathobionts. Nature Communications 12, 6664–6680 (2021).

26. Sprotte, S., Brinks, E., Neve, H. & Franz, C.M. Complete genome sequence of the novel virulent phage PMBT24 infecting Enterocloster bolteae from the human gut. Heliyon 10, e28813 (2024).

27. Ferrocino, I. & Pérez, M.R. Gut-microbiotatargeted diets modulate human immune status. Revista de nutrición práctica, 19–19 (2022).

28. Cani, P.D., Depommier, C., Derrien, M., Everard, A. & de Vos, W.M. Akkermansia muciniphila: paradigm for next-generation beneficial microorganisms. Nature Reviews Gastroenterology & Hepatology 19, 625–637 (2022).

29. Buswell, J.A., Odier, E. & Kirk, T.K. Lignin biodegradation. Critical Reviews in Biotechnology 6, 1–60 (1987).

30. Cui, L., et al. Lignin biodegradation and its valorization. Fermentation 8, 366–384 (2022).

31. Sood, A., et al. The probiotic preparation, VSL#3 induces remission in patients with mild-to-moderately active ulcerative colitis. Clinical Gastroenterology and Hepatology 7, 1202–1209 (2009).

32. Chapman, T.M., Plosker, G.L. & Figgitt, D.P. VSL#3 probiotic mixture: a review of its use in chronic inflammatory bowel diseases. Drugs 66, 1371–1387 (2006).

33. Chen, K. et al. High internal phase emulsions stabilized with carboxymethylated lignin for encapsulation and protection of environmental sensitive natural extract. International journal of biological macromolecules 158, 430–442 (2020).

34. Wang, J. et al. Reduction of lignin color via one-step UV irradiation. Green Chemistry 18, 695–699 (2016).

35. Dong, X. et al. Antimicrobial and antioxidant activities of lignin from residue of corn stover to ethanol production. Industrial Crops and Products 34, 1629–1634 (2011).

36. Yang, Y.Q. et al. Amphiphilic copolymer brush with random pH-sensitive/hydrophobic structure: synthesis and self-assembled micelles for sustained drug delivery. Soft Matter 8, 454–464 (2012).

37. Cheng, C. et al. Qing-Chang-Hua-Shi granule ameliorates DSS-induced colitis by activating NLRP6 signaling and regulating Th17/Treg balance. Phytomedicine 107, 154452–154463 (2022).

38. Zhang, C. et al. Comprehensive analysis of the characteristics and differences in adult and newborn brown adipose tissue. Diabetes 67, 1759–1769 (2018).

39. Elmentaite, R. et al. Cells of the human intestinal tract mapped across space and time. Nature 597, 250–255 (2021).

40. Lee, M. et al. Protein stabilization of ITF2 by NF-κB prevents colitis-associated cancer development. Nature Communications 14, 2363–2380 (2023).

