## Supplemental Information for "Orally surface engineered probiotic system for ulcerative colitis therapy *via* modulating gut microbiota and immune homeostasis"

**Material and Methods**

**The derivation process of mathematical expressions related to the plug flow model.**

**For the release process**

$r_A = 0.601\ C_{CML@EcN0} - 0.0873$ (Fig.2L)

$r_A = 0.601\ (1\text{-}X) - 0.0873 = -0.601X+0.5137$

$-r_A = F_{A0}\, dX/dV$

$dF = -r_A dV$, $vdC = -r_A vdt$, $dC = -r_A dt = -r_A(C)\ dt$

∵ $C = 1\text{-}X$

∴ $dX = r_A dt = r_A(X)\ dt$

$r_A$: Reaction rate（g/mL min）

F: Flow（g/min）

C: Initial concentration（g/mL）

V: Volum（mL）

v: Material movement speed（mL/min）

Thus:

$dX = (-0.601X+0.5137)\ dt$

$$\frac{dX}{-0.601X+0.5137} = dt$$

Integrating both sides simultaneously yields:

$$\frac{ln\ (-0.601X+0.5137)}{-0.601} = t + C_1 \qquad (1)$$

When t = 0, X = 0.

Bring to (1), $C_1 \approx 1.11$

$$\text{Namely: } X = \frac{e^{[-0.601\ (t + 1.11)] - 0.5137}}{-0.601} \qquad (2)$$

Bring X = 0.5 to (2), $t_{1/2} \approx 1.46$ h

Bring t = 20 to (2), X ≈ 0.86 (86% of *EcN* release)

**For the proliferation process**

$r_A = 0.0135\ C_{EcN0} + 0.0202$ (Fig.2K)

$r_A = 0.0135\ (1\text{-}X) + 0.0202 = -0.0135X + 0.0337$

Similarly:

$dX = (-0.0135X + 0.0337)\ dt$

$$\frac{dX}{-0.0135X + 0.0337} = \mathrm{dt}$$

Integrating both sides simultaneously yields:

$$(\ln(-0.0135X + 0.0337))/(-0.0135) = t + C_2 \quad (3)$$

When t = 0, X = 0.

Bring to (1), $C_2 \approx 251.22$

Namely: $$X = \frac{e^{[-0.0135\,(t + 251.22)] - 0.0337}}{-0.0135} \quad (4)$$

Add (2) and (4) together:

$$X = \frac{e^{[-0.601\,(t + 1.11)] - 0.5137}}{-0.601} + \frac{e^{[-0.0135\,(t + 251.22)] - 0.0337}}{-0.0135} \quad (5)$$

X: Conversion rate

t: Reaction time

Bring t = 20 to (5), X ≈ 0.86 + 0.59 = 1.45 (86% came from CML@*EcN* Release, 59% from *EcN* proliferation. In the total *EcN* population, 14% exist in the form of CML@*EcN*, while 131% are present as free *EcN* colonizing the colon).

The initial gavage volume is 2 * 10 ^ 8 CFU. According to the model calculation, the final settlement *EcN* is approximately:

$5*1.31*2*10^8 = 1.31*10^9$ CFU (gavage for 5 days in treatment experiment), which is approximately 0.1% of the total GM of mice.

**Supplementary Table 1 |** Optimization of reaction conditions

| Samples | Lignin (g) | $ClCH_2COONa$ (g） | Temperature (°C) |
|---|---|---|---|
| CML1 | 7.0492 | 1.5 | 85 |
| CML2 | 7.0492 | 4.5 | 85 |
| CML3 | 7.0492 | 7.5 | 85 |

**Supplementary Table 2 |** Parameters of CMLs

| Samples | Ph-OH (mmol/g) | -COOH (mmol/g) | Z (mV) |
|---|---|---|---|
| Lignin | 2.7085 | 1.6889 | -16.74 |
| CML1 | 2.7632 | 2.2118 | -23.55 |
| CML2 | 2.3179 | 2.4064 | -25.16 |
| CML3 | 2.2556 | 3.2439 | -31.83 |

**Supplementary Table 3 |** Flow cytometry antibody information

| Manufacturer | Lot number | Antibodies and Color Matching |
|---|---|---|
| BioLegend | 103105 | PE anti-mouse CD45 |
| BioLegend | 103106 | PE anti-mouse CD45 |
| BioLegend | 101211 | APC anti-mouse/human CD11b |
| BioLegend | 101212 | APC anti-mouse/human CD11b |
| BioLegend | 115511 | APC anti-mouse CD19 |
| BioLegend | 115512 | APC anti-mouse CD19 |
| BioLegend | 127627 | Brilliant Violet 421™ anti-mouse Ly-6G |
| BioLegend | 127628 | Brilliant Violet 421™ anti-mouse Ly-6G |
| BioLegend | 123137 | Brilliant Violet 421™ anti-mouse F4/80 |
| BioLegend | 123131 | Brilliant Violet 421™ anti-mouse F4/80 |
| BioLegend | 123132 | Brilliant Violet 421™ anti-mouse F4/80 |
| BioLegend | 117329 | Brilliant Violet 421™ anti-mouse CD11c |
| BioLegend | 117343 | Brilliant Violet 421™ anti-mouse CD11c |
| BioLegend | 117330 | Brilliant Violet 421™ anti-mouse CD11c |
| BioLegend | 137612 | Brilliant Violet 421™ anti-mouse CD335 (NKp46) |
| BioLegend | 137611 | Brilliant Violet 421™ anti-mouse CD335 (NKp46) |

**Supplementary Table 4 |** Primer sequences

| Primer | Forward (5‘–3’) | Reverse (5‘–3’) |
|---|---|---|
| V3V4a | GGACTACHVGGGTWTCTAAT | ACTCCTACGGGAGGCAGCA |
| fim | ATACTACGACGGTAAATGGT | TACATCAGTATCGGTAGCAT |
| foc | CCACGGTTAGGTGTGGTACA | CGTCGGCGTTGGCAATACCA |
| pMUT2a | GACCAAGCGATAACCGGATG | GTGAGATGATGGCCACGATT |
| ZO-1 | GGCAAAAACAGAAGGATTGC | TAAGCCGGCTGAGATCTTGT |
| Occludin | ACAGCTTTCTGGGTGGATT | TGAGGACCGCTAGCAAGTTT |
| Cludin-1 | GTCCATAGGCACCGTATTGC | CCCATGCTGGAAAAACACTT |
| GAPDH | AGGTTGTCTCCTGCGACTTCA | TGGTCCAGGGTTTCTTACTCC |
| ZO-1 | GCAAGGGATAGAAGCGCAAG | TATGGCTGGCCAATCGAAGAC |
| Occludin | GGACTACNNGGGTATCTAAT | CGGTCCATCTTTCTTCGGGT |
| Claudin-1 | CATCAATGCCAGGTATGAATT | TGTTGGGTAAGAGGTTGTTT |

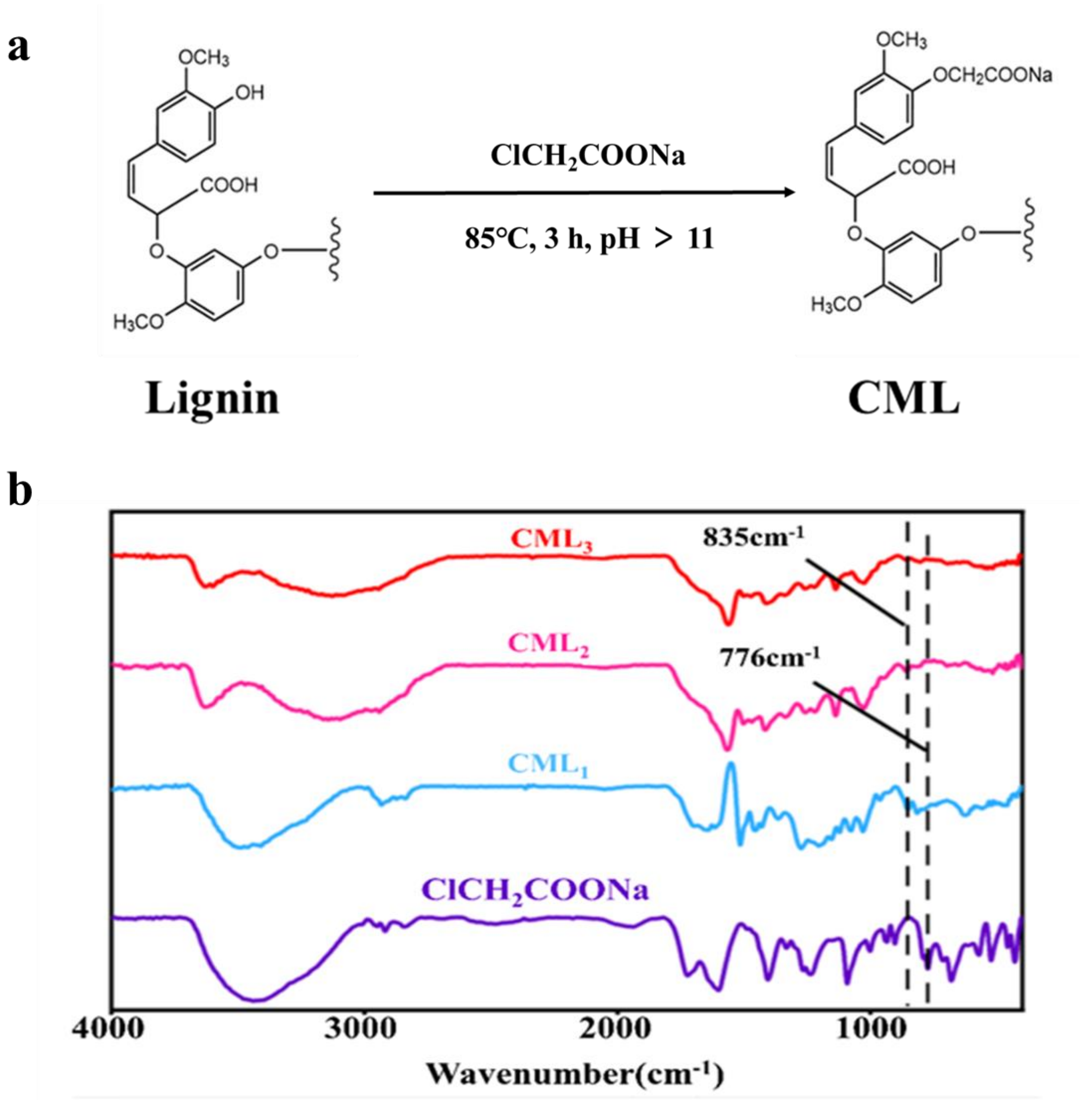


**Supplementary Fig. 1.** (a) Reaction formula of lignin modification. (b) FTIR differential spectrum of CML and unmodified lignin was used to verify the success of lignin modification.

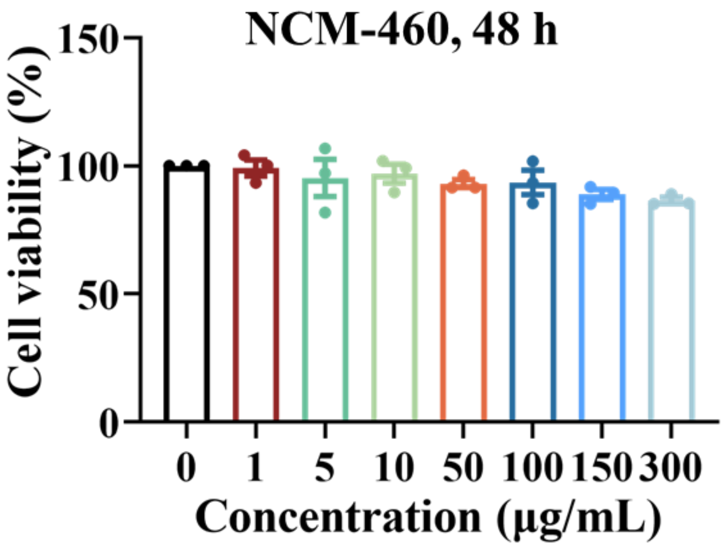


**Supplementary Fig. 2.** Cytotoxicity of CML against NCM-460 cell with incubation for 48 hours.

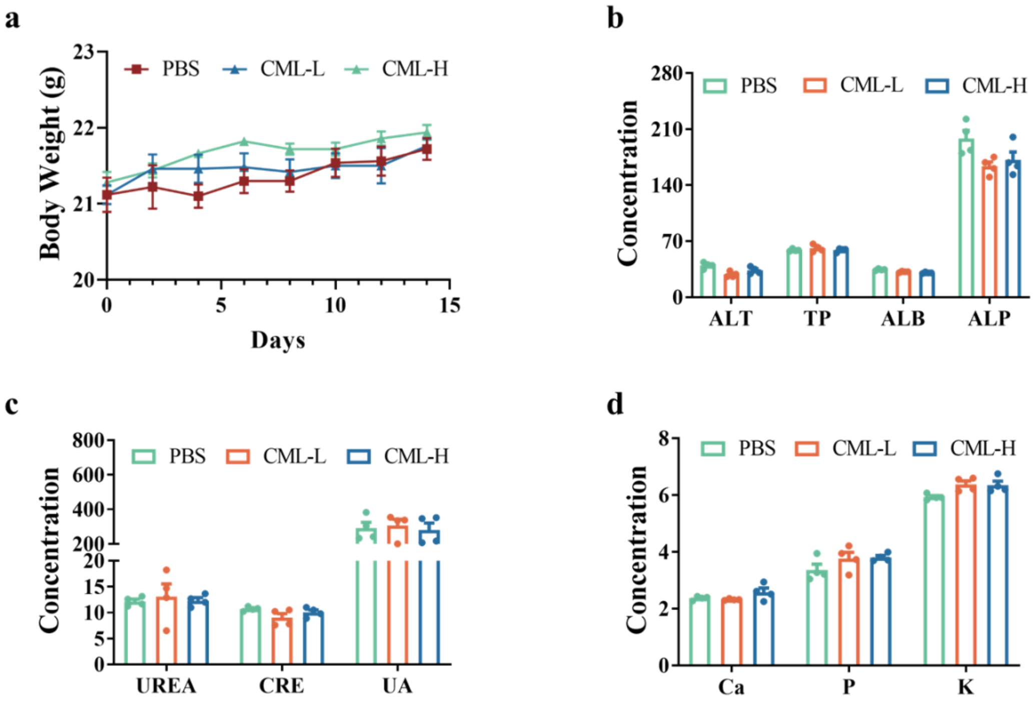


**Supplementary Fig. 3. *In vivo* biosafety validation of CML.** 15 days of continuous intervention of CML did not significantly affect the (a) body weight and blood biochemical indicators for (b) liver function, (c) kidney function, (d) inorganic ion of mice.

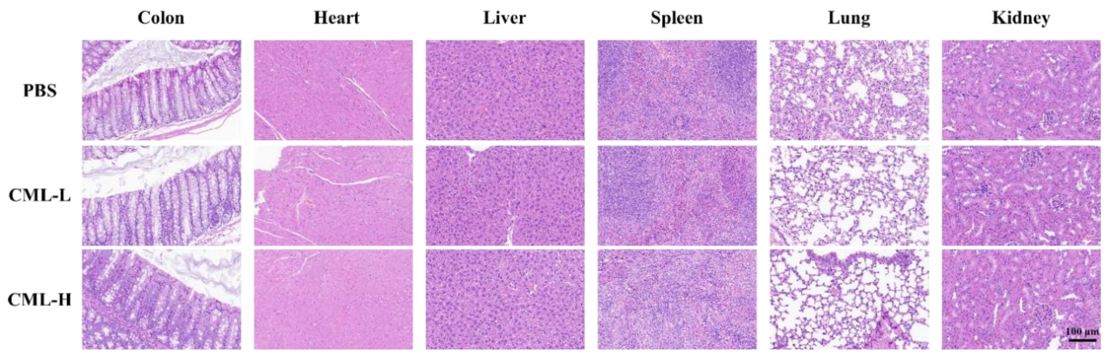


**Supplementary Fig. 4. *In vivo* biosafety validation of CML.** 15 days of continuous intervention of CML did not significantly affect the morphological characteristics of important organ tissues of mice. Scale bar, 100 μm.

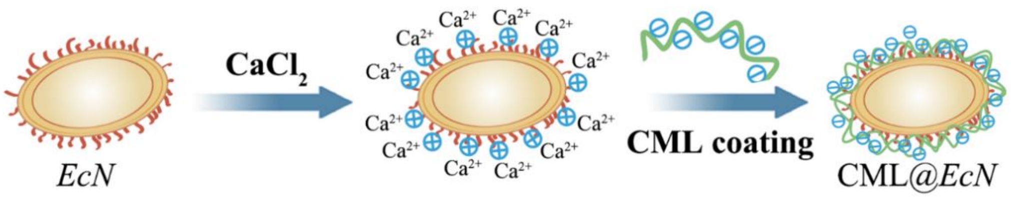


**Supplementary Fig. 5.** Diagram of CML@*EcN* preparation process.

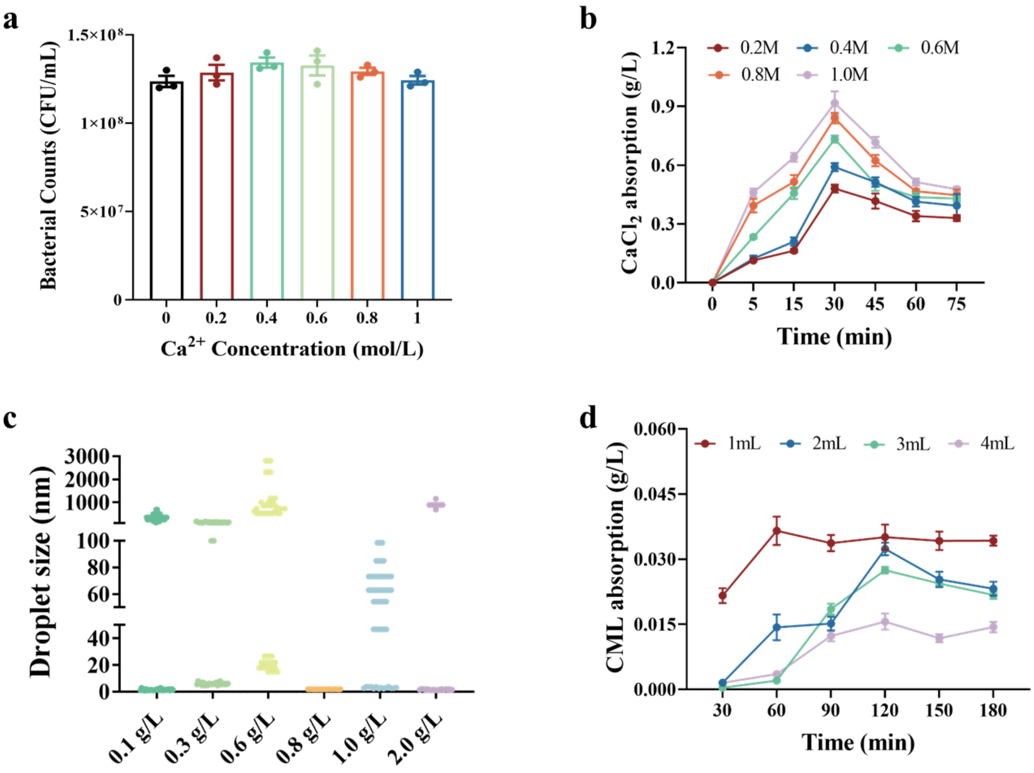


**Supplementary Fig. 6.** Optimization of (a-b) $Ca^{2+}$ concentration and (c-d) CML concentration during the preparation process of CML@*EcN*.

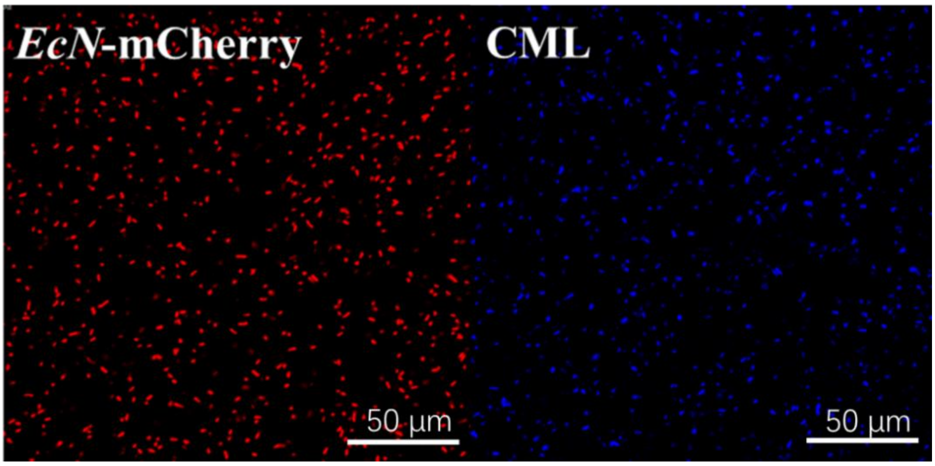


**Supplementary Fig. 7.** Single channel confocal image of *EcN* and CML. Scale bar, 50 μm.

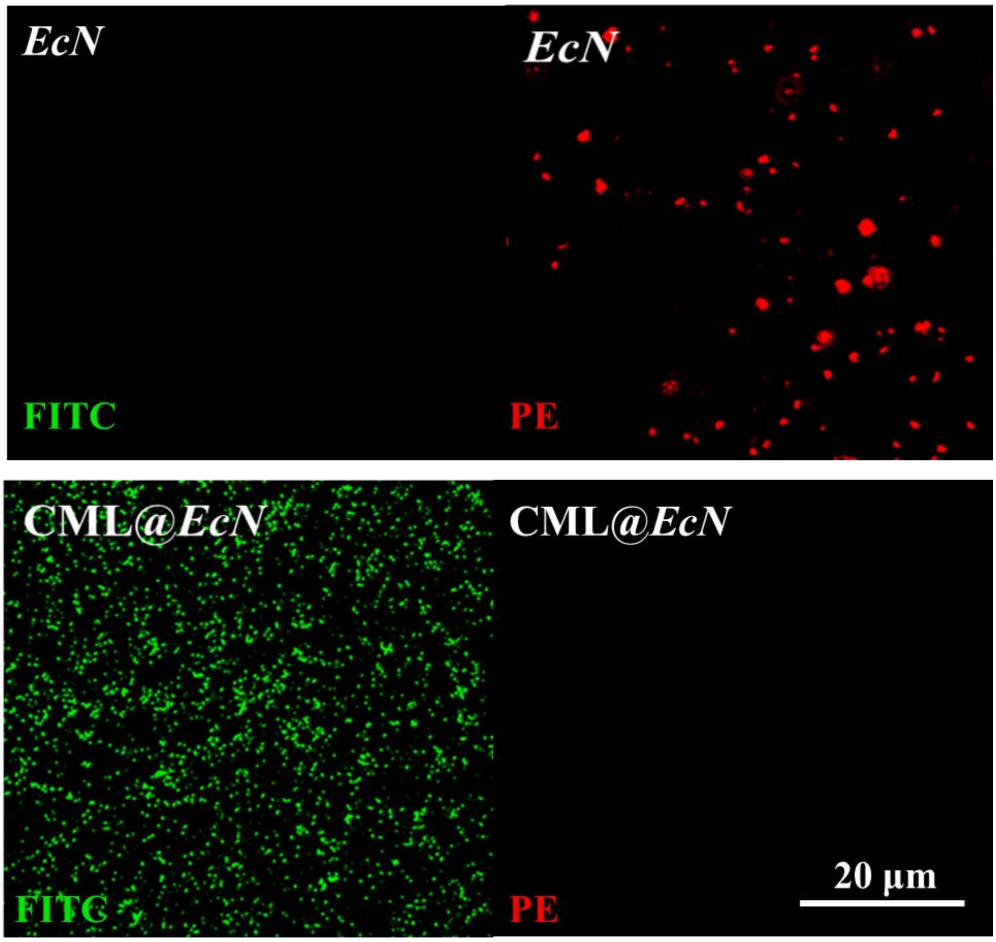


**Supplementary Fig. 8.** Single channel image of *in vitro* simulated microenvironment experiment. Scale bar, 20 µm.

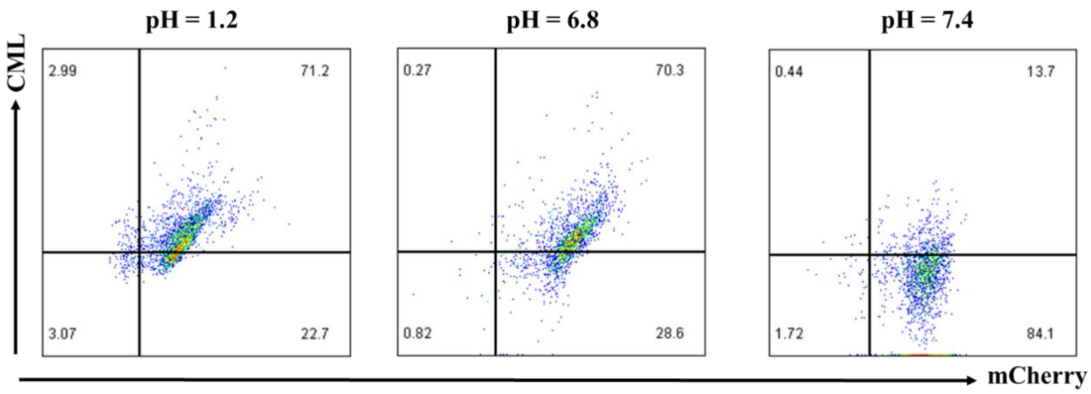


**Supplementary Fig. 9. CML@*EcN* has pH responsiveness.** After 8 hours of incubation, *EcN* was almost not released in the environment of pH = 1.2 and pH = 6.8, and was largely released in the alkaline environment of pH = 7.4.

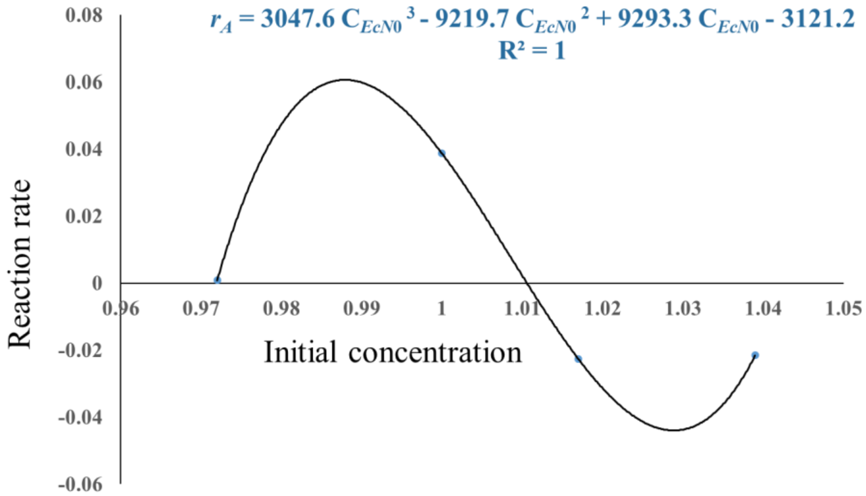


**Supplementary Fig. 10. Fitting curve of CML@*EcN* in the small intestine.**

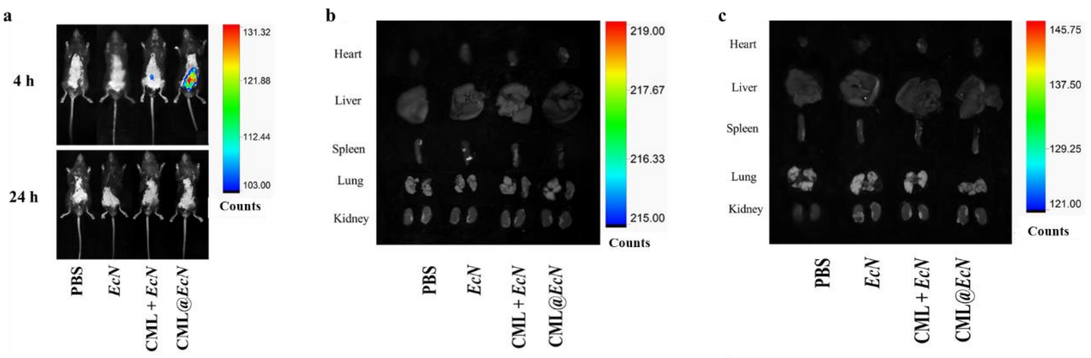


**Supplementary Fig. 11.** Qualitative distribution of *EcN in vivo* at 4 and 24 hours (a) mice and (b-c) important organs after gavage, respectively.

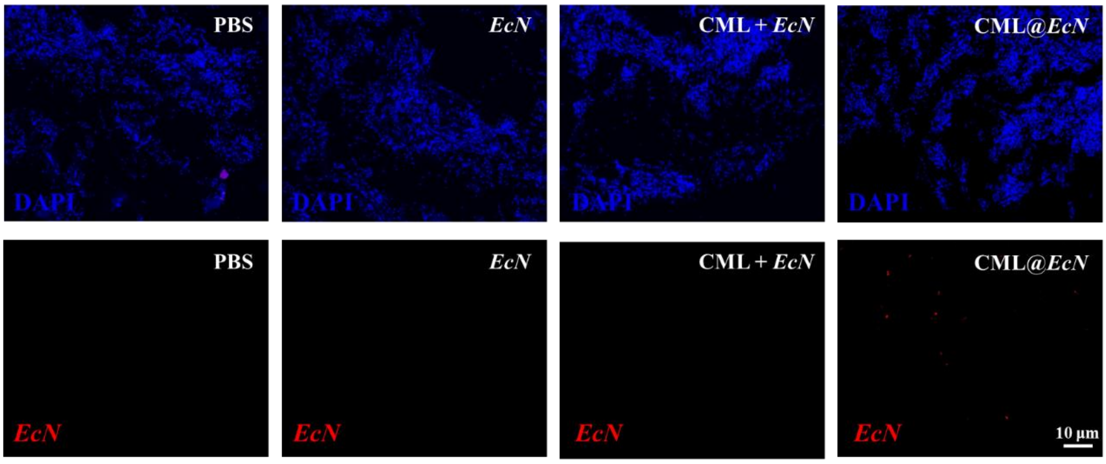


**Supplementary Fig. 12.** Single channel FISH image of DAPI and *EcN*. Scale bar, 10 μm.

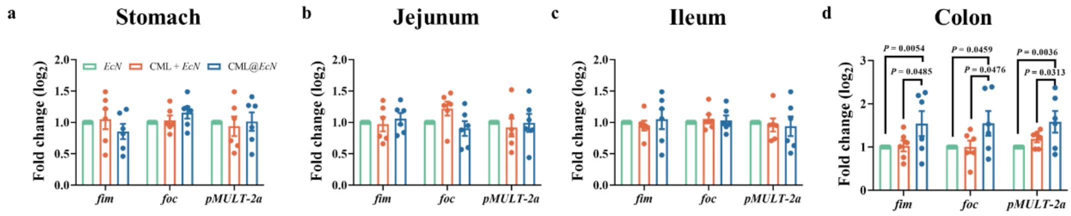


**Supplementary Fig. 13.** Quantitative distribution of *EcN in vivo* at 72 hours.

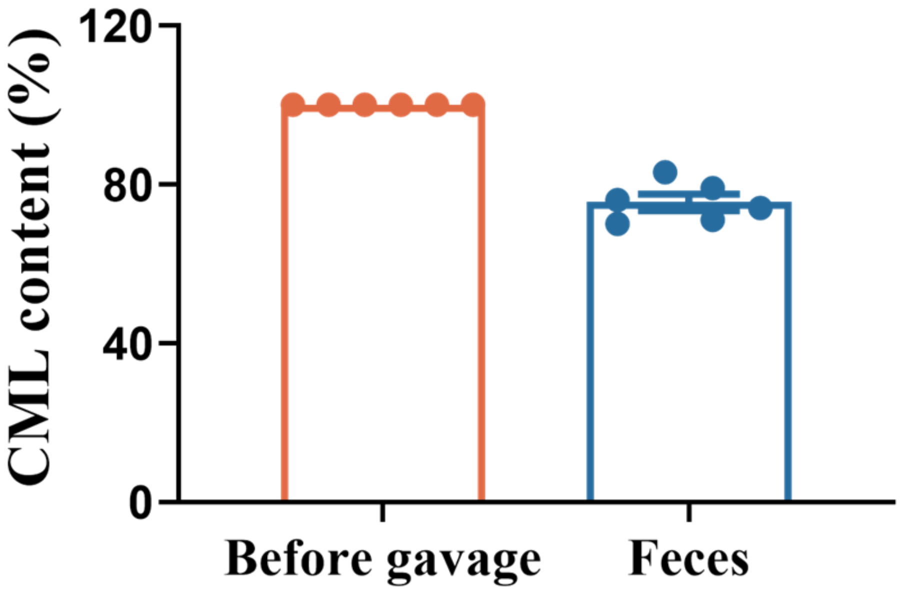


**Supplementary Fig. 14.** Proportion of CML excreted with faeces at 20 hours. (n = 6, n represents independent experiments).

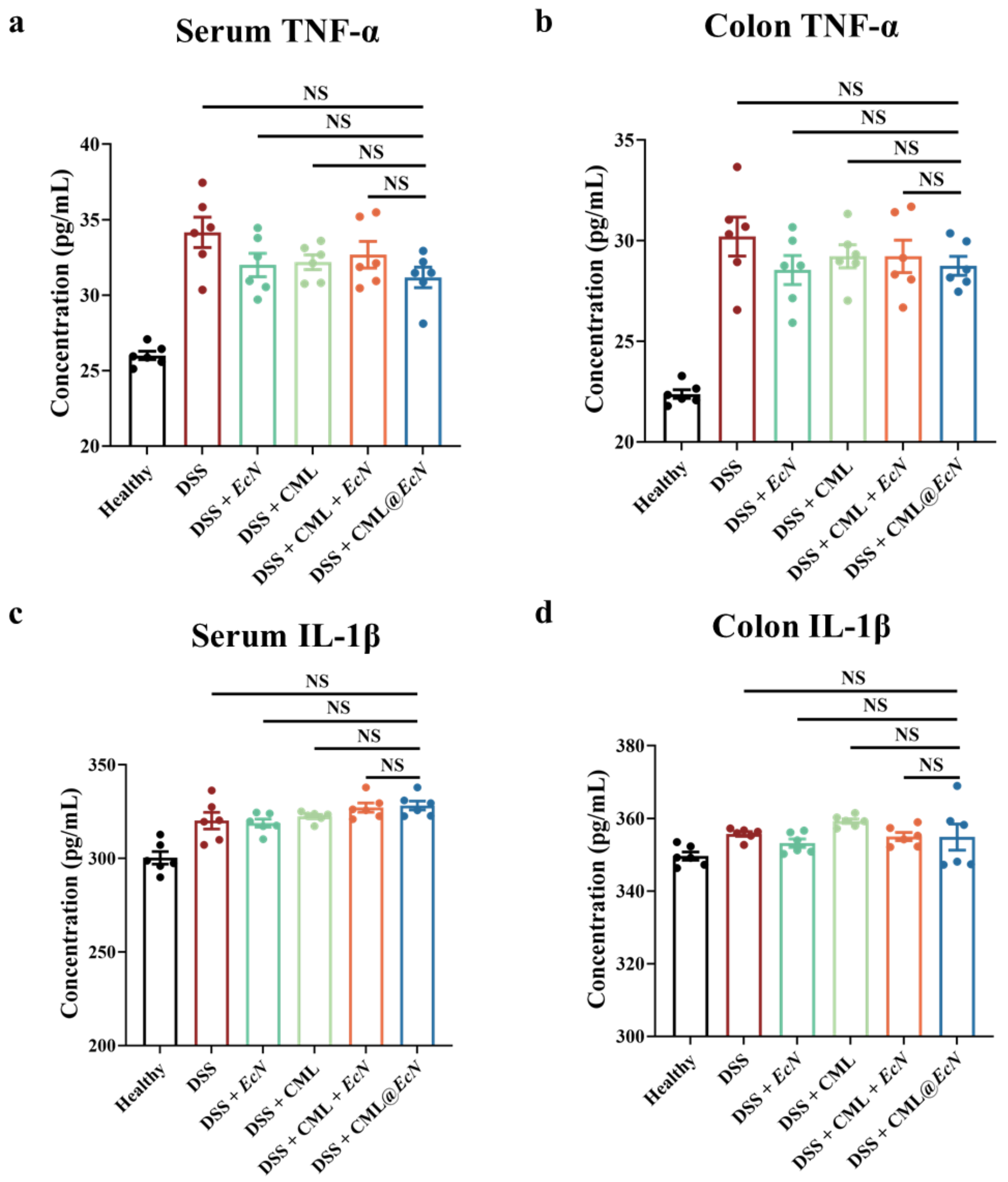


**Supplementary Fig. 15. The levels of inflammatory factors in the serum of mice in each group.** After 7 days of intervention, serum and colon samples from each group of mice were used for the detection of inflammatory factors. The levels of (a-b) TNF-*α* and (c-d) IL-1*β* in the CML@*EcN* group showed no significant difference compared to the DSS and other intervention groups. (n = 6, n represents independent experiments).

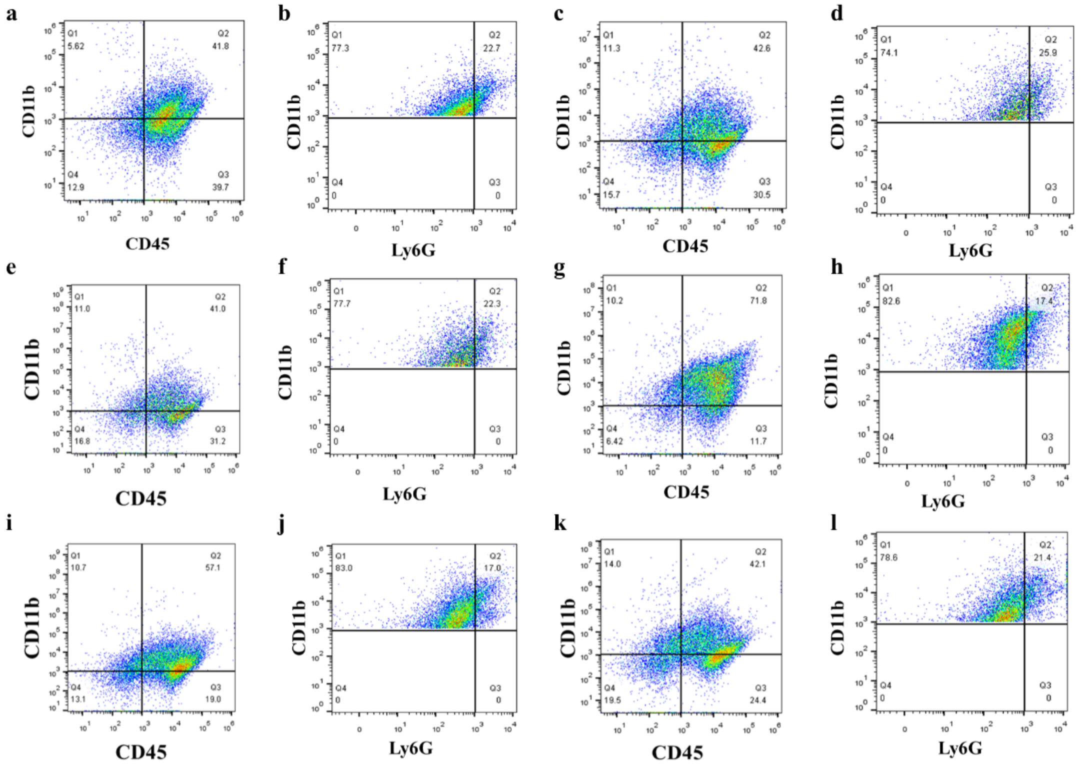


**Supplementary Fig. 16.** Flow cytometry classifying of neutrophils in the colon of mice in (a-b) Healthy, (c-d) DSS, (e-f) DSS + *EcN*, (g-h) DSS + CML, (i-j) DSS + CML + *EcN* and (k-l) DSS + CML@*EcN* group.

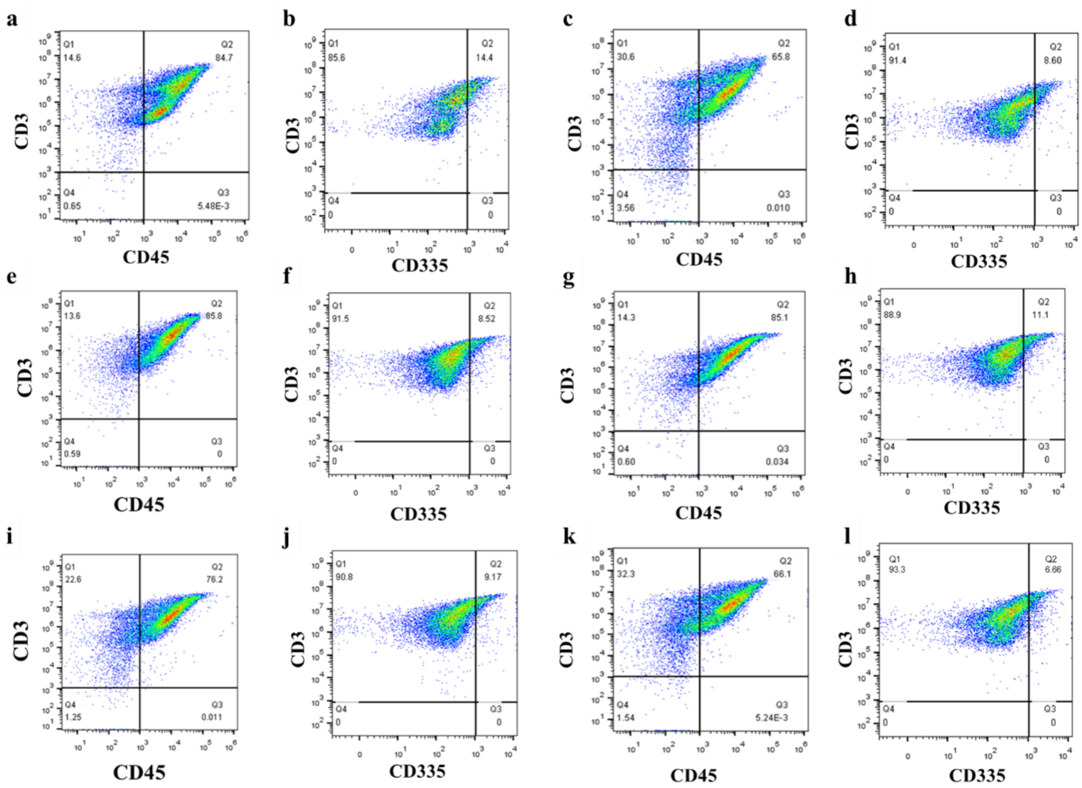

**Supplementary Fig. 17.** Flow cytometry classifying of T cells in the colon of mice in (a-b) Healthy, (c-d) DSS, (e-f) DSS + *EcN*, (g-h) DSS + CML, (i-j) DSS + CML + *EcN* and (k-l) DSS + CML@*EcN* group.

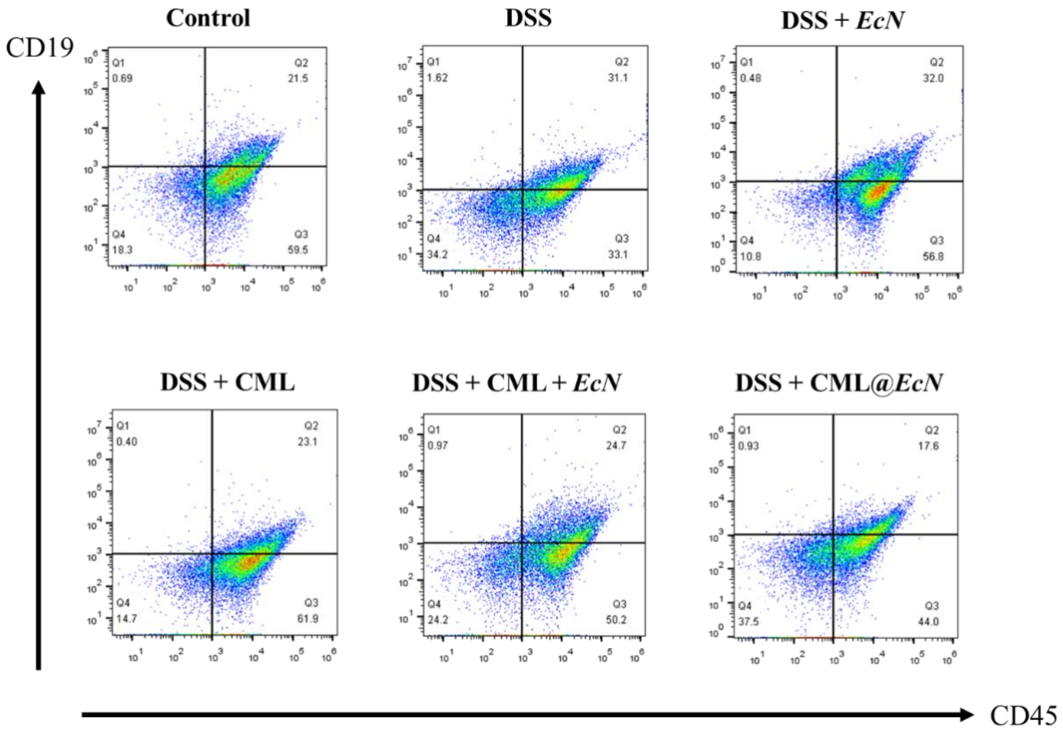


**Supplementary Fig. 18.** Flow cytometry classifying of B cells in the colon of mice in each group.

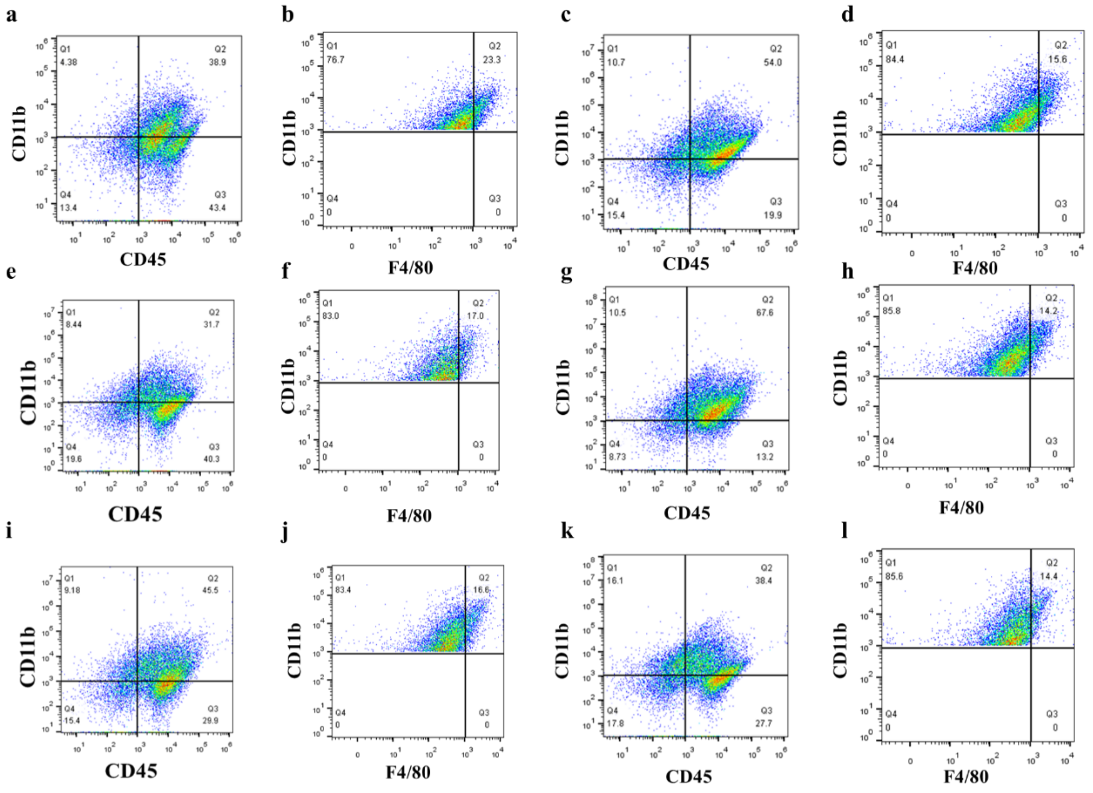


**Supplementary Fig. 19.** Flow cytometry classifying of macrophages in the colon of mice in (a-b) Healthy, (c-d) DSS, (e-f) DSS + *EcN*, (g-h) DSS + CML, (i-j) DSS + CML + *EcN* and (k-l) DSS + CML@*EcN* group.

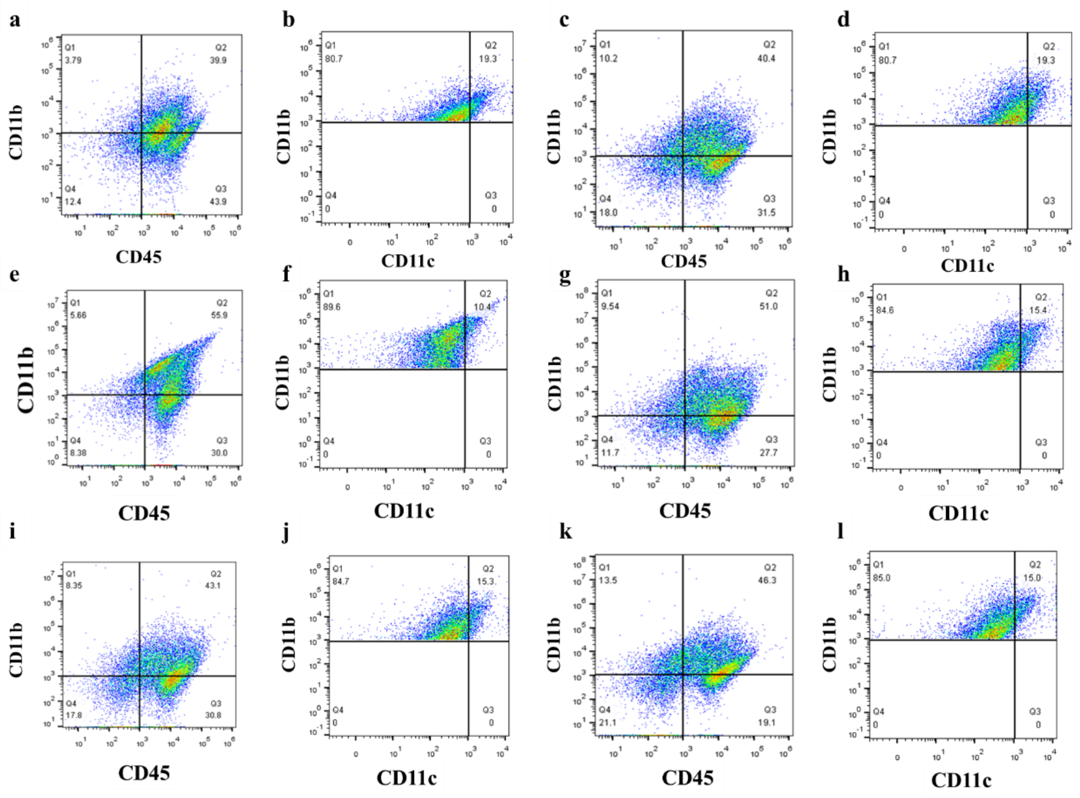


**Supplementary Fig. 20.** Flow cytometry classifying of DCs in the colon of mice in (a-b) Healthy, (c-d) DSS, (e-f) DSS + *EcN*, (g-h) DSS + CML, (i-j) DSS + CML + *EcN* and (k-l) DSS + CML@*EcN* group.

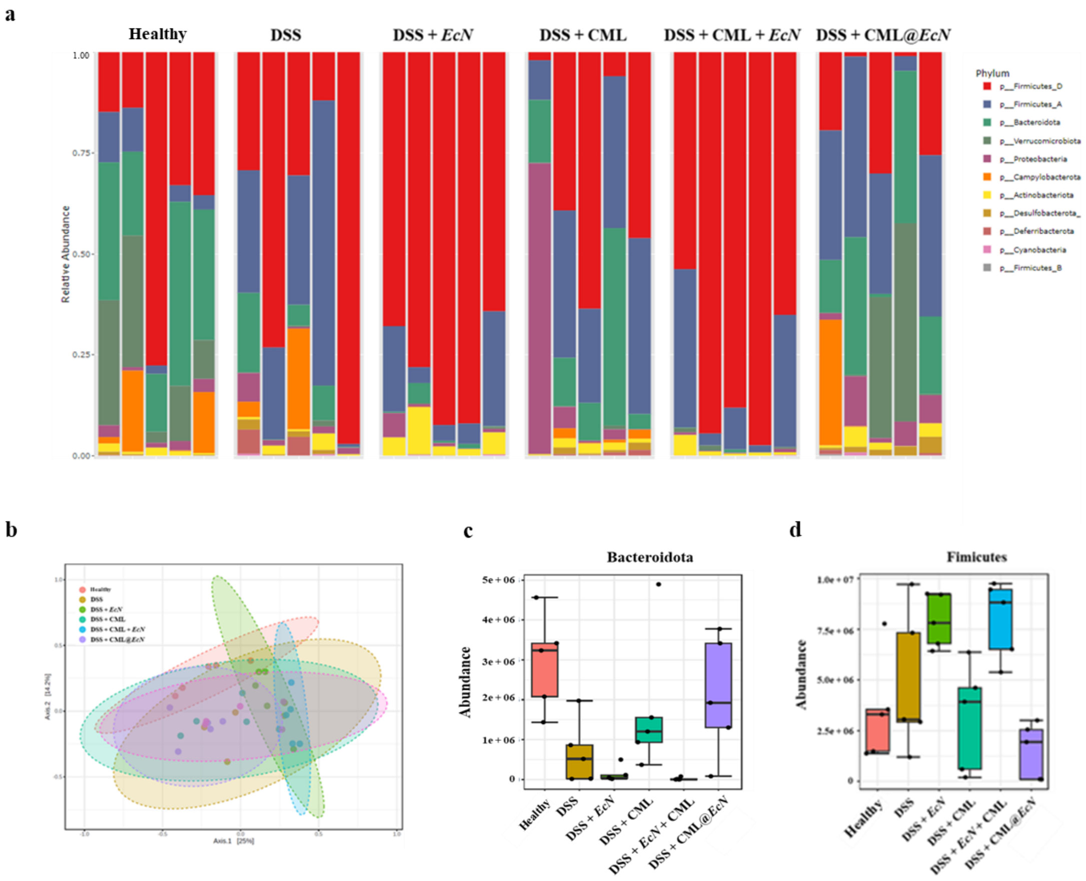


**Supplementary Fig. 21.** (a-b) The overall structure of GM in each group of mice. (c-d) The two phylum with the largest differences in GM.

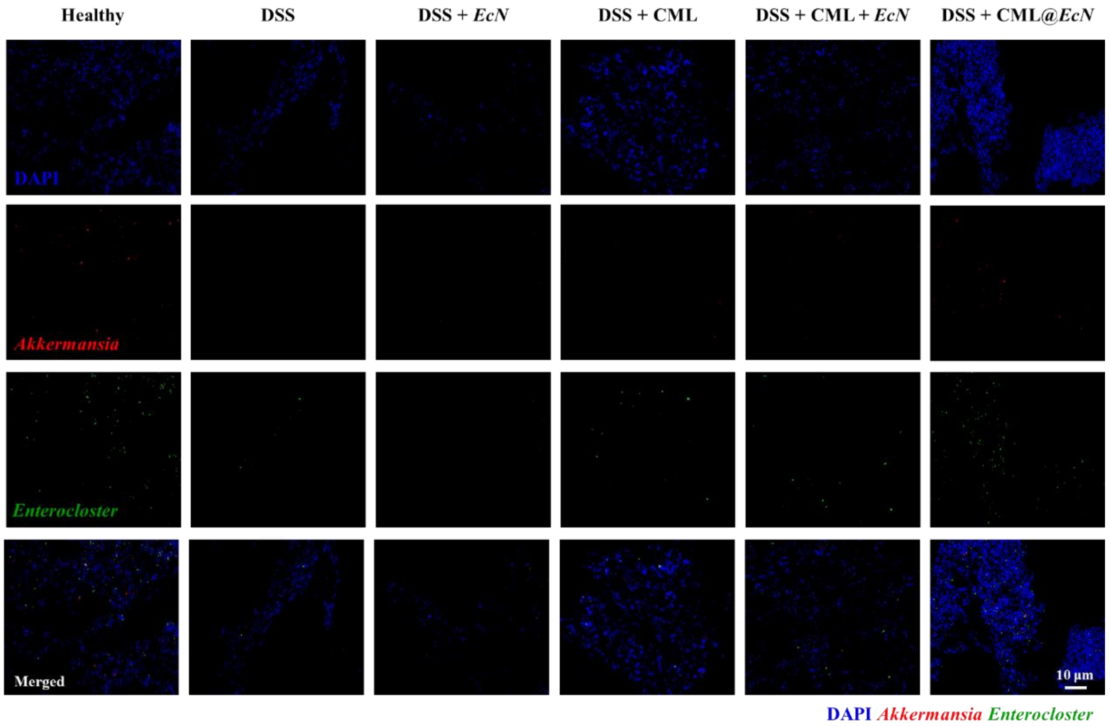


**Supplementary Fig. 22.** Single channel image of double probe labeled FISH for *Akkermansia* & *Enterocloster*. Scale bar, 10 μm.

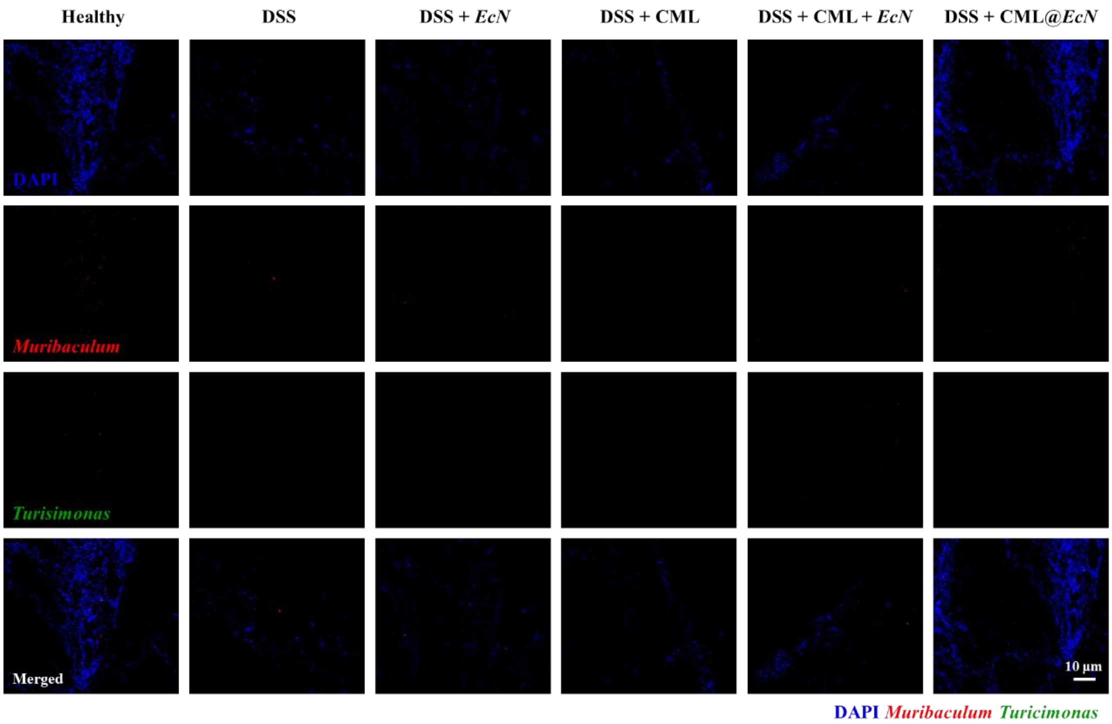


**Supplementary Fig. 23.** Single channel image of double probe labeled FISH for *Muribaculum* & *Turisimonas*. Scale bar, 10 μm.

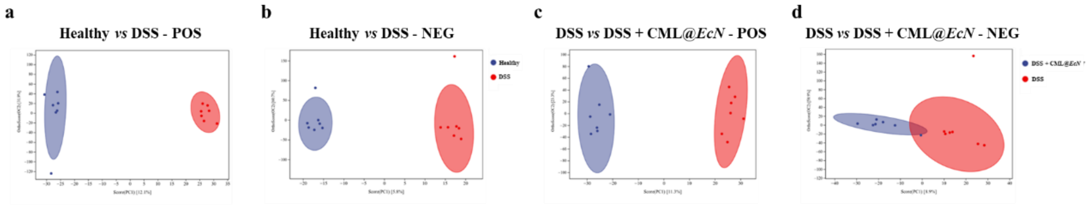


**Supplementary Fig. 24.** OPLS-DA analysis of differential metabolites in the cationic and anionic mode.
